# An Accessible and Scalable Pipette-Driven Microfluidic Platform for Lipid Nanoparticle Formulation

**DOI:** 10.64898/2026.11.06.757988

**Authors:** Yuji S. Takeda, Fujun Wang, Xin Gu, James Forster, Haoyi Hou, Parnian Hemmati, Behnam Partopour, Laura Maugeri, Yuriy Kim, Sabrina Apel, Ryan Norton, Joseph Finelli, Parsa Molaei, Shayan Shafiee, Sixuan Li, Jinghan Lin, Chen-Wei Chiang, Martha Mayo, Seyoum Ayehunie, Samin Akbari, Hai-Quan Mao, Tza-Huei Wang, Mehdi Dehghani

**Affiliations:** Sartorius Stedim North America Inc., Marlborough, MA 01752; MatTek, Now Part of Sartorius, Ashland, MA 01721; Department of Mechanical Engineering Johns Hopkins University, Baltimore, MD 21218; Institute for NanoBioTechnology, Johns Hopkins University, Baltimore, MD 21218; CICS Analytics Corp., Baltimore, MD 21218; Department of Materials Science and Engineering, Johns Hopkins University, Baltimore, MD, 21218; Department of Biomedical Engineering, Johns Hopkins University School of Medicine, Baltimore, MD 21218; Translational Therapeutic and Regenerative Engineering Center, Johns Hopkins University School of Medicine, Baltimore, MD 21218, USA

**Keywords:** Lipid nanoparticle, Microfluidic mixer, High throughput screening, mRNA delivery, Scale-up manufacturing, Single-particle analysis

## Abstract

Advances in lipid nanoparticle (LNP) technology have increased the need for accessible and scalable methods that accelerate low-volume formulation screening. Here, we describe a disposable microfluidic mixer actuated by a positive-displacement pipette equipped with dual-chamber pipette tip, providing volumetric control at a fixed flow rate ratio without external microfluidic pumps. Computational fluid dynamics analysis predicted shorter micromixing times in the high-shear spiral channel than in the low-shear design. Across three benchmark lipid formulations, the high-shear mixer reproducibly generated smaller LNPs with higher mRNA encapsulation efficiency than the low-shear mixer or manual pipetting. Orthogonal characterization further revealed higher particle concentrations, structural features consistent with compact particles, more uniform mRNA loading, and a lower empty-particle fraction. LNPs prepared using the selected high-shear mixer mediated functional mRNA delivery in 2D cell cultures, a 3D human liver tissue model, and mice. Integration with a liquid-handling robot enabled automated screening of formulation parameters. Dimensionless mixing-time analysis guided enlargement of the mixer geometry for a separate production system, which produced 75 mL preparations at 150 mL/min with particle size and mRNA encapsulation efficiency comparable to those obtained with the small-scale mixer. This platform provides an accessible, automation-compatible approach to LNP formulation screening and a route to larger-scale production.

---

Lipid nanoparticles (LNPs) are an established delivery platform for mRNA therapeutics, with applications expanding beyond infectious-disease vaccines to targeted extrahepatic delivery, cancer vaccines, chimeric antigen receptor (CAR) T therapies, and gene editing.^1–8^ In addition to lipid composition and RNA design, mixing conditions during preparation can influence particle size, RNA encapsulation, empty particle fraction, and functional cargo delivery.^9,10^ Mixing governs the rate and spatial uniformity of ethanol dilution as the lipid and aqueous streams combine, thereby shaping the local environment in which LNP assembly occurs.^11–13^

Reliable comparison of LNP compositions during formulation screening requires small-volume preparation methods that minimize variability arising from mixing. Commercial LNP mixers, such as NanoAssemblr Spark (Cytiva) and Sunscreen (Unchained Labs), support small-volume LNP preparation, but their capital cost can limit access for smaller research laboratories, including academic groups and startups. Lab-built microfluidic systems can offer a lower-cost alternative, although optimizing channel geometry and achieving precise flow control remain technical challenges for reproducible production of small, uniform particles.^14^

Liquid-handling robots offer a route for high-throughput screening (HTS) through automated pipetting, but conventional pipette-based LNP preparation offers limited control over microscale mixing and can yield larger particles than microfluidic mixing.^15–19^ This motivates the development of accessible and disposable microfluidic mixers that provide controlled mixing in both manually operated and automated formulation workflows.

Here, we introduce an accessible, disposable, pipette-driven microfluidic mixer for LNP preparation (Fig. 1a, S1). The device couples a dual-chamber pipette tip to a microfluidic mixer and is actuated by an electronic positive-displacement pipette, providing volumetric control at a fixed flow rate ratio without external microfluidic pumps. We evaluated mixing performance using computational fluid dynamics (CFD), compared the resulting LNPs using orthogonal analytical methods, and assessed functional mRNA delivery by LNPs prepared using the selected high-shear design in HEK293T cells, a 3D human liver tissue model, and mice. Integration with a liquid-handling robot enabled automated screening of formulation parameters. Finally, dimensionless mixing-time analysis guided scale-up of the mixer geometry for use in a separate production system. Larger-volume preparations yielded LNPs with particle size and mRNA encapsulation efficiency comparable to those obtained with the small-scale mixer.

**Figure 1.**
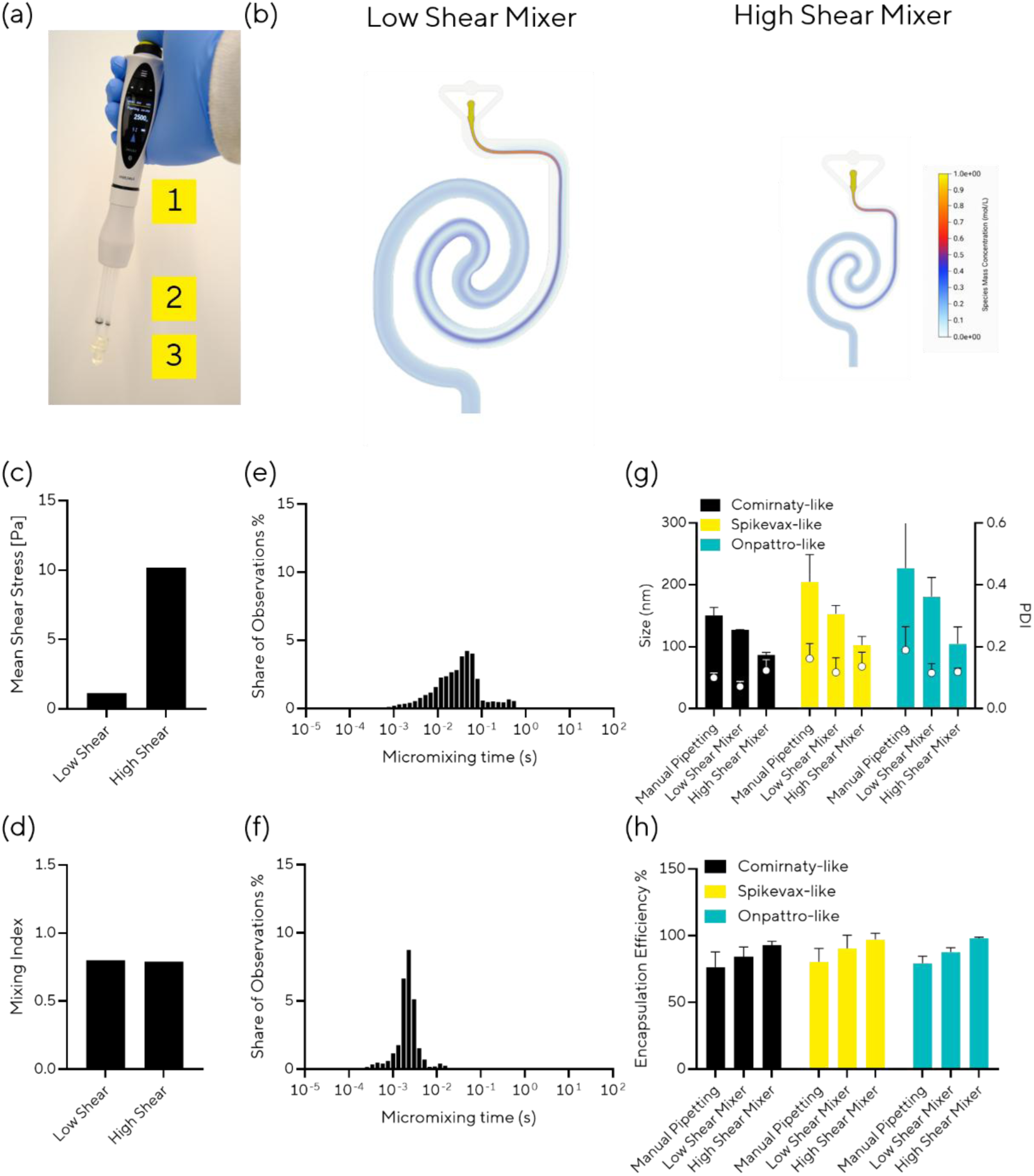
Development of a pipette-driven mixer for lipid nanoparticle preparation. (a) Overview of the device components: (1) Picus® 2 electronic pipette, (2) dual-chamber pipette tip, and (3) microfluidic mixer chip. (b–f) CFD analysis of mixing performance. (b) Steady-state passive-scalar concentration profiles in the low-shear and high-shear mixers. (c) Mean shear stress across the entire channel. (d) Mixing index at each channel outlet. (e, f) Distributions of micromixing times calculated for individual computational cells throughout the (e) low-shear and (f) high-shear mixer channels. (g) Particle size and polydispersity index (PDI) measured by dynamic light scattering (DLS). (h) mRNA encapsulation measured by the RiboGreen assay. Size and RNA encapsulation data are presented as mean ± standard deviation with n = 3 independent measurements.

## Results and Discussion

### Pipette Mixer Design and Channel Geometry Selection

Commercial LNP formulation systems use costly, sophisticated equipment for precise flow control, whereas lab-built setups require calibrated external pumps and tubing that introduce dead volume. To simplify this workflow, we developed a single-use, pipette-driven LNP formulation tool comprising a dual-chamber pipette tip (component 2 in Fig. 1a) coupled to a microfluidic mixer (component 3 in Fig. 1a). The tip’s chambers has a volumetric ratio of 3:1, providing precise volumetric control and a fixed 3:1 flow rate ratio of aqueous mRNA solution to lipid solution. This design enabling LNP formulation through a standard pipetting workflow without dedicated pumping equipment (Fig. S1).

The spiral-shaped mixer was selected because spiral microchannels generate Dean-vortex secondary flows, which enhance cross-sectional mass transfer under laminar flow conditions. Previous studies have shown that spiral micromixers achieve high mixing efficiency by promoting rapid homogenization of fluid streams.^20,21^ CFD analysis was used to distinguish three aspects of mixing: shear stress characterizes hydrodynamic stresses associated with local velocity gradients; the outlet mixing index quantifies the spatial uniformity of the simulated concentration field; and micromixing time estimates the local timescale for concentration homogenization. These metrics are complementary rather than interchangeable: similar outlet mixing indices do not necessarily indicate comparable local mixing times within the channel. The two channel geometries generated mean shear stresses of 1.1 and 10.2 Pa and were therefore designated low-shear and high-shear, respectively (Fig. 1c). Despite this difference, the low- and high-shear geometries had similar mixing indices (0.80 and 0.79, respectively) (Fig. 1d). We therefore additionally evaluated the predicted micromixing-time distributions to distinguish local mixing conditions that were not captured by the outlet mixing index.

Rapid mixing plays a critical role in nanoparticle nucleation, with recent studies suggesting that micromixing, rather than macroscopic mixing, is especially important for LNP formation.^11–13^ Micromixing time characterizes the timescale over which local concentration gradients are reduced through the combined effects of fluid deformation and molecular diffusion. This timescale influences LNP size by determining how many nuclei form before the lipid monomer pool is depleted. Rapid and homogeneous micromixing promotes a synchronized burst of supersaturation, generating many nuclei that each incorporate a smaller fraction of available lipid and therefore form smaller, more uniform particles; Khan *et al.* reported smaller lipid nanoparticles at approximately 20 ms than at 60–100 ms under matched lipid concentrations and flow rate ratios.^11^ Slower local ethanol dilution can produce fewer nuclei that continue to grow, resulting in larger particles with broader size distributions, as proposed by Maeki *et al.*, who emphasized that complete mixing was not required for particle-size control.^13^ CFD-based estimates indicated shorter micromixing times in the high-shear channel (mean: 2.54 ms; median: 2.12 ms) than in the low-shear channel (mean: 71.5 ms; median: 32.2 ms). The high-shear design also exhibited narrower distributions, as indicated by its lower coefficient of quartile variation (CQV) values, compared with the low-shear design (Fig. 1e, f, Table S1). The shorter predicted micromixing times and narrower micromixing-time distribution led us to postulate that a high-shear geometry, which combines rapid and homogeneous mixing, would provide particularly favorable conditions for LNP assembly.

To test the predicted effects of micromixing, we experimentally evaluated two channel designs using EGFP mRNA-loaded LNPs with lipid compositions based on those of Comirnaty, Spikevax, and Onpattro (Table 1). Before encapsulation, the purity of in-house synthesized mRNA was assessed by PATfix analytical chromatography using the LNP Switcher and the IVT Monitoring modules.^22,23^ The LNP Switcher analysis confirmed mRNA purity after purification, while the IVT Monitoring module, designed for at-line monitoring of mRNA synthesis and NTP consumption, provided greater resolution for assessing the removal of residual NTPs (Fig. S2).

**Table 1.** LNP formulations tested in this study. The percentages in the table represent the molar ratio of each lipid component.

|  | **Comirnaty-like** | **Spikevax-like** | **Onpattro-like** |
|---|---|---|---|
| **Ionizable lipid** | 46.3% ALC-0315 | 50% SM-102 | 50% MC3 |
| **Phospholipid** | 9.4% DSPC | 10% DSPC | 10% DSPC |
| **Cholesterol** | 42.7% | 38.5% | 38.5% |
| **PEGylated lipid** | 1.6% ALC-0159 | 1.5% DMG-PEG2000 | 1.5% DMG-PEG2000 |

We then evaluated particle size, polydispersity index (PDI), and RNA encapsulation. As a control, LNPs were prepared by 10 cycles of manual pipetting, a typical workflow using a liquid-handling for HTS.^15–19^ Across all three formulations, the high-shear mixer produced LNPs with smaller particle sizes and lower PDI, measured by dynamic light scattering (DLS), and higher RNA encapsulation efficiency, measured by RiboGreen assay, than the low-shear mixer (Fig. 1g, h). The manual pipetting control generated larger LNPs than high- and low-shear mixer conditions and resulted in the lowest RNA encapsulation efficiency. The PDI of all mixer-prepared samples was below 0.15, indicating relatively narrow particle size distribution. Together, these results suggest that particle size, PDI, and encapsulation efficiency were not fully explained by the mixing index, which reflects macroscopic-scale concentration homogenization, and that micromixing time may play a more important role in governing particle formation. These experimental findings support the CFD-based prediction that the high-shear mixer provides more favorable conditions for producing small, uniform LNPs with high encapsulation efficiency.

DLS and RiboGreen assay provide critical information on particle size and bulk RNA encapsulation, respectively. Recent studies have highlighted the value of high-resolution and orthogonal analytical methods for revealing additional biophysical and biochemical characteristics of LNPs, including morphology, particle-to-particle compositional variability, RNA payload distributions, and empty-particle fractions.^24–28^ Here, properties of LNPs produced by the pipette-driven mixers were further examined using additional analytical tools.

First, particle concentration and size were evaluated using nanoparticle tracking analysis (NTA) in scattering (Sc-NTA) and fluorescence modes (Fl-NTA), which has been used to measure the size and concentration of lipid-based nanoparticles and extracellular vesicles.^24^ LNPs produced with the high-shear mixer showed higher particle concentrations (4.4–4.8 × 10^12^ particles/mL) than those produced with the low-shear mixer (1.5–3.8 × 10^12^ particles/mL) or manual pipetting (1.4–2.4 × 10^12^ particles/mL) (Fig. 2a, b). No statistically significant differences in particle concentration were detected between Sc-NTA and Fl-NTA; the corresponding particle size distributions are shown in Fig. S3.

**Figure 2.**
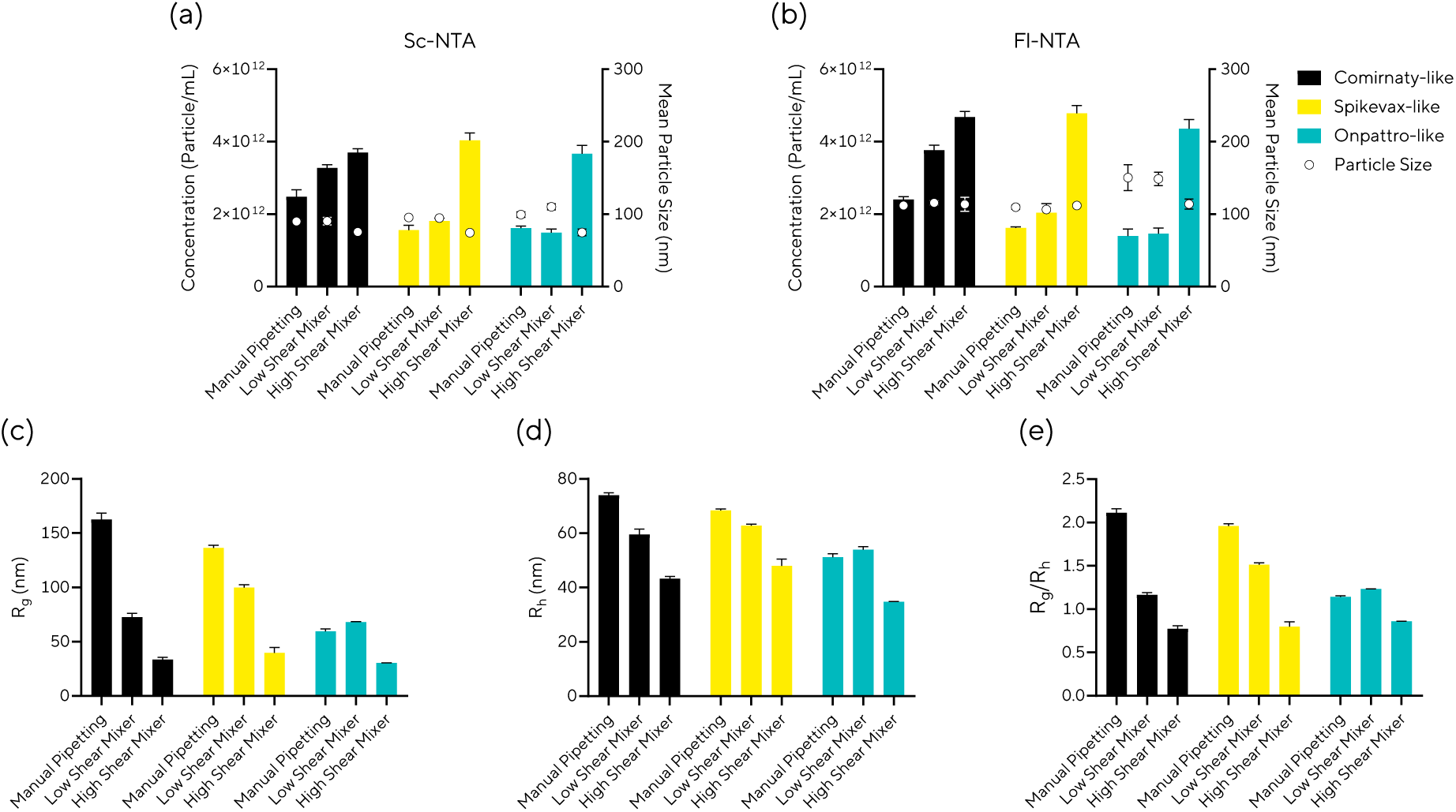
LNP concentration and structural characterization. (a, b) Particle concentration and size determined by nanoparticle tracking analysis in (a) scattering (sc-NTA) and (b) fluorescence (fl-NTA) modes. (c–e) Asymmetric field-flow fractionation coupled with MALS/QELS (AF4-MALS/QELS) analysis was used to determine (c) radius of gyration (R_g_), (d) hydrodynamic radius (R_h_), and (e) shape factor (R_g_/R_h_). All data are presented as mean ± standard deviation with n = 3 independent measurements.

Structural characteristics were subsequently assessed using asymmetric-flow field-flow fractionation coupled with multi-angle light scattering and quasi-elastic light scattering (AF4–MALS/QELS), which provides indirect morphological information beyond particle size alone.^25^ MALS and QELS measured the radius of gyration (R_g_) and hydrodynamic radius (R_h_), respectively, and to calculate the shape factor, ρ = R_g_/R_h_, as an indicator of shape-related structural properties (Fig. 2c–e, S4). LNPs produced with the high-shear mixer eluted earlier and had smaller R_h_ and R_g_ values than the low-shear mixer and manual pipetting. LNPs produced with the high-shear mixer had shape factors of 0.77, 0.80, and 0.86 for the Comirnaty-like, Spikevax-like, and Onpattro-like formulations, respectively. These values were close to the theoretical value of 0.775 for a homogeneous solid sphere and therefore indicative of relatively dense particle structures.^26^ In the same formulation order, shape factors were higher for the low-shear mixer (1.2, 1.5, and 1.5) and manual pipetting (2.1, 2.0, and 1.1), exceeding the theoretical benchmarks for homogeneous solid spheres (0.775) and thin hollow spheres (1.0).

We next examined whether mixing geometry affects single-particle homogeneity in lipid composition and cargo loading. Single-Particle Automated Raman Trapping Analysis (SPARTA) enables detection and identification of individual lipid components within single LNPs.^27^ Using SPARTA, we analyzed Raman fingerprints in three spectral regions corresponding to the ionizable lipids—ALC-0315 (1655–1677 cm⁻¹), SM-102 (1120–1145 cm⁻¹), and MC3 (1639–1673 cm⁻¹)— as well as cholesterol (696–714 cm⁻¹) and total lipids (1415–1500 cm⁻¹) (Fig. 3, Table S2–S10). To assess particle-to-particle compositional homogeneity, CQV values were calculated for each Raman fingerprint and compared across formulations. LNPs prepared with the high-shear mixer tended to exhibit lower CQV values for ionizable lipid (0.087, 0.16, and 0.034 for the Comirnaty-like, Spikevax-like, and Onpattro-like formulations, respectively) than manual pipetting (0.21, 0.13, and 0.052) and the low-shear mixer (0.17, 0.28, and 0.03), indicating improved ionizable lipid homogeneity across individual particles. However, the high-shear mixer did not consistently yield the lowest lipid CQV values across formulations and lipid components; for example, SM-102 CQV was lower with manual pipetting than with the high-shear mixer in the Spikevax-like formulation. Together, these results suggest that the effect of mixing conditions on particle-to-particle lipid homogeneity depends on the formulation and lipid component.

**Figure 3.**
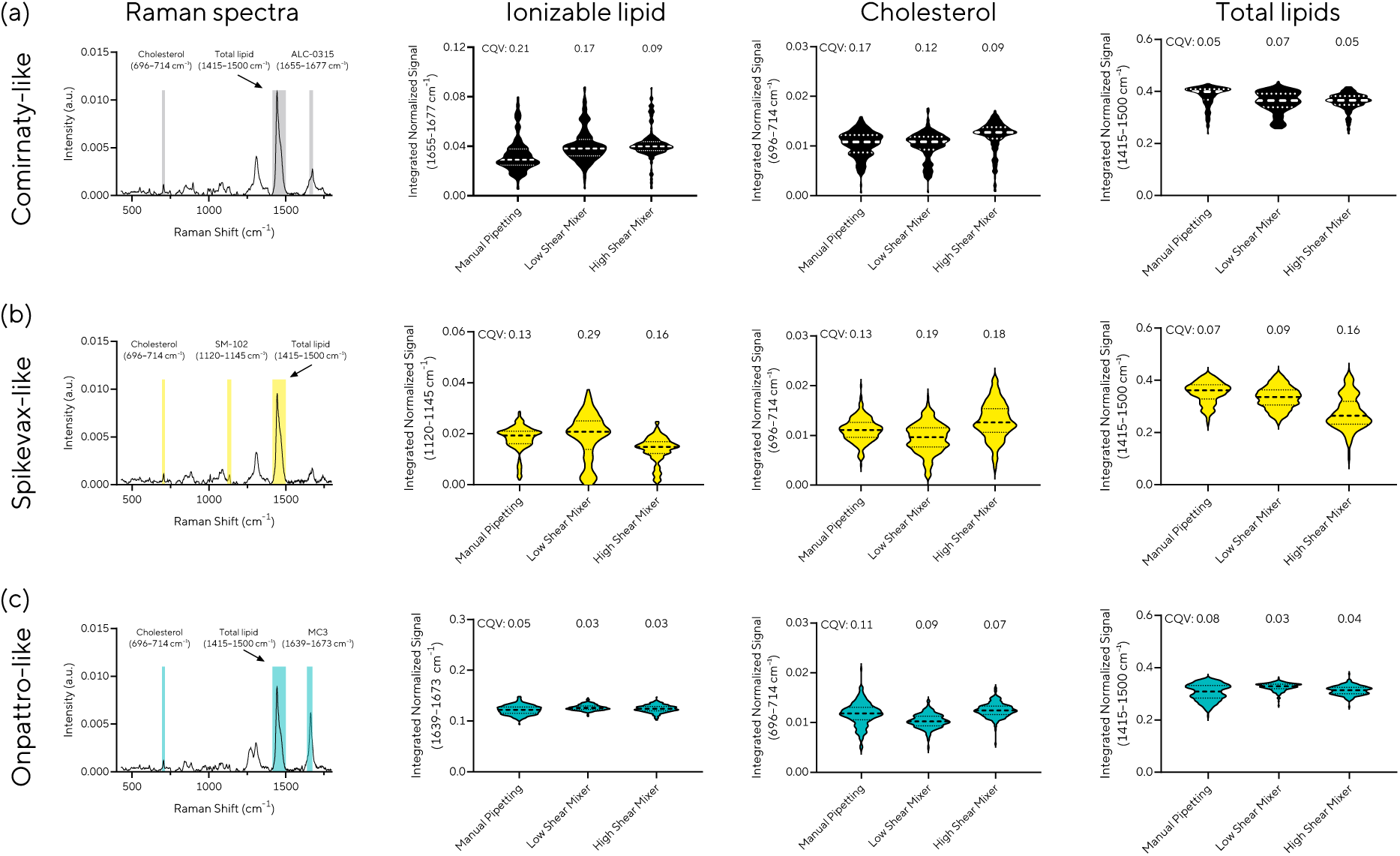
SPARTA characterization of LNPs prepared using different mixer geometries. The left columns present mean Raman spectra for (a) Comirnaty-like, (b) Spikevax-like, and (c) Onpattro-like formulations. The three right columns display violin plots of normalized lipid Raman intensity across individual LNPs, indicating relative lipid content and particle-to-particle compositional variability. Dashed and dotted lines mark the median and the first and third quartiles (Q1 and Q3), respectively. Each plot reports the coefficient of quartile variation (CQV) for each sample. Approximately 250 particles were analyzed per sample.

To complement the SPARTA analysis of lipid compositional variability, we next performed multi-color cylindrical illumination confocal spectroscopy (CICS) to assess mRNA loading heterogeneity in individual LNPs (Fig. 4, S5). CICS is a flow-based, multicolor single-particle fluorescence technique that distinguishes mRNA-loaded LNPs, empty LNPs, and free mRNA through fluorescence coincidence analysis. By deconvolving particle fluorescence against single-mRNA signals, CICS quantifies the mRNA copy-number distribution and payload capacity of individual LNPs.^29–31^ For CICS analysis, LNPs were prepared using cyanine 5 (Cy5)–labeled mRNA encoding EGFP and Top Fluor-PC (0.5 mol% of total lipid) to label the mRNA and lipid components, respectively. Labeling did not appreciably alter particle size. The high-shear mixer produced a lower fraction of empty LNPs (37–40%) and higher encapsulation efficiency (86–88%) across all three formulations than the low-shear mixer (42–56% empty LNPs; 46–66% encapsulation) and manual pipetting (61–67% empty LNPs; 38–56% encapsulation) (Fig. 4a, b). Although LNPs prepared with the high-shear mixer contained fewer mRNA copies per particle (6.1–7.3 copies/particle), their mRNA copy-number distribution was more uniform than those obtained with the low-shear mixer (8.7–10.4 copies/particle) or manual pipetting (6.6–8.8 copies/particle), both of which produced broader, more heterogeneous distribution (Fig. 4c, d, Table S11–S13). These findings highlight the importance of evaluating both payload distribution and empty-particle fraction. Truong *et al.* reported that RNA payload per LNP and the fraction of empty particles can influence gene-editing performance *in vivo*.^28^ Together, the SPARTA and CICS data support the CFD-based prediction that the high-shear mixer promotes more homogeneous mixing.

**Figure 4.**
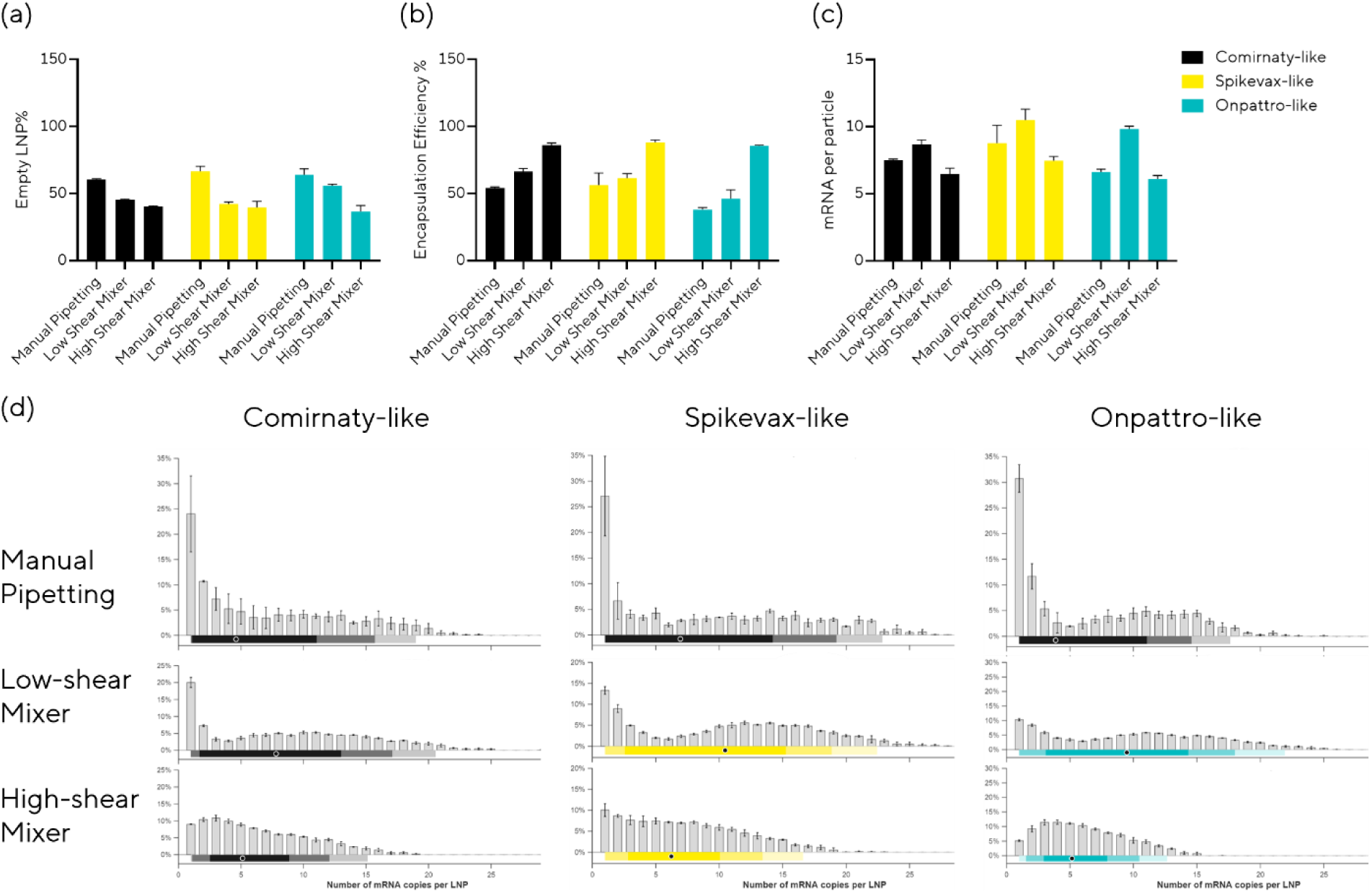
LNP population analysis across mixer geometries and shear conditions using cylindrical illumination confocal spectroscopy (CICS). (a) Percentage of empty nanoparticles. (b) Encapsulation efficiency measured by CICS. (c) Mean mRNA copy number per particle. (d) mRNA payload distribution per particle. Dots beneath each histogram indicate the median. All data are presented as mean ± standard deviation with n = 3 independent measurements.

Taken together, the high-shear mixer produced smaller and more uniform particles than the low-shear mixer and manual pipetting control. Across the orthogonal analytical methods, these LNPs consistently exhibited compact structures and more uniform mRNA loading. Based on these results, the high-shear mixer geometry was selected for subsequent studies.

### Functional Evaluation and High-Throughput Formulation Screening

We first assessed the morphology of LNPs produced with the high-shear pipette-driven mixer by cryo-EM. Representative images showed predominantly spherical particles (Fig. 5a). In order to evaluate the functionality of LNPs formulated with the high shear mixer, we then evaluated the transfection activity of EGFP mRNA-loaded LNPs in HEK293T cells using time-lapse fluorescent imaging system. EGFP fluorescence was detected within 12 h, increased with mRNA dose, and persisted for at least 48 h after dosing (Fig. 5b–e, S6). Live-cell counts remained comparable between treated and untreated groups (Fig. S6d–f).

**Figure 5.**
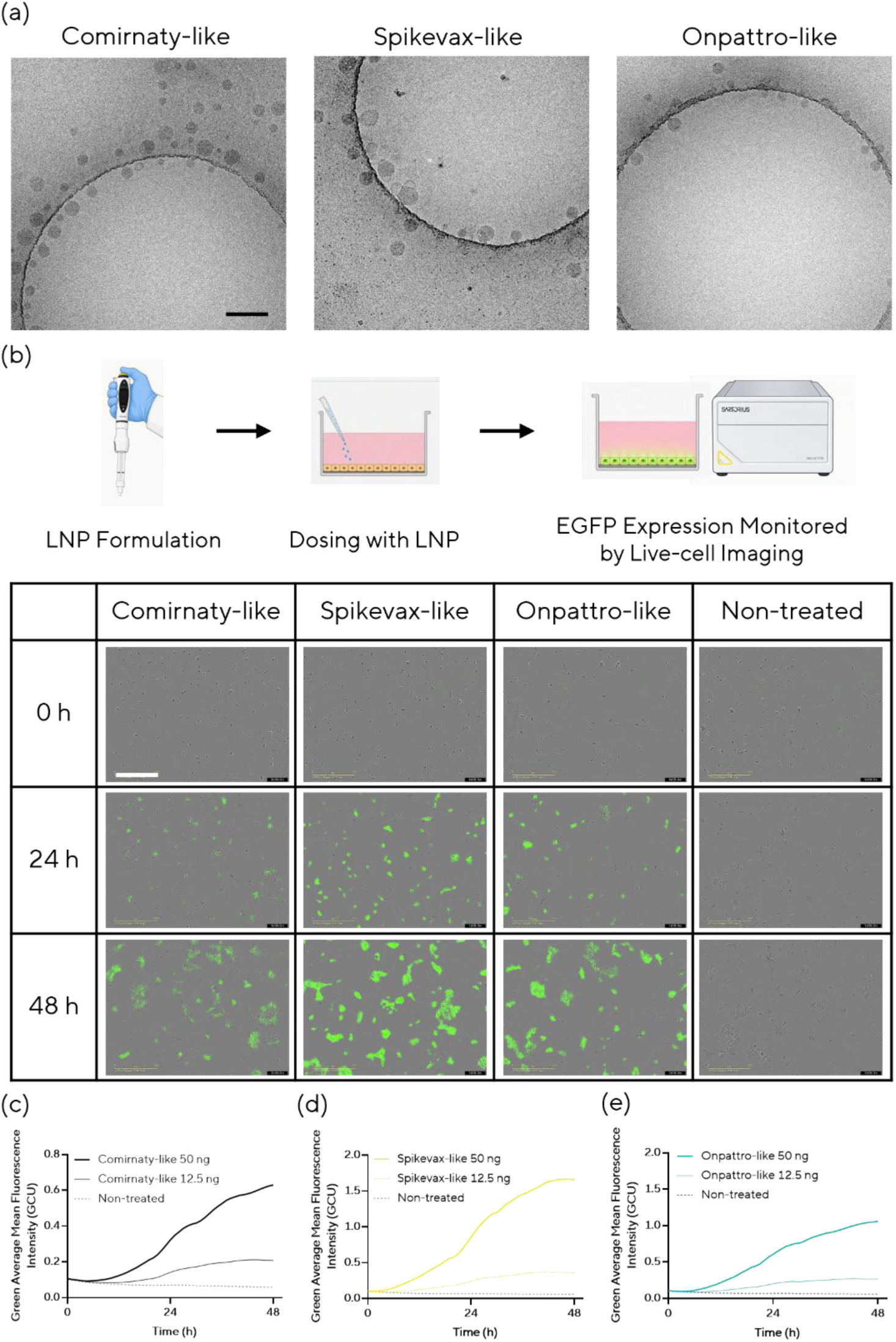
LNP morphology and transfection activity in HEK293T cells. (a) Representative cryo-EM images of LNPs prepared with the high-shear mixer. Scale bar, 200 nm. (b–e) Functional evaluation of LNPs in 2D cell culture using an *in vitro* transfection assay. HEK293T cells were dosed with LNPs prepared using the high-shear mixer, and transfection was monitored using the IncuCyte live-cell imaging system. Each sample was tested in three wells, with two images acquired per well at each time point. Illustrations in the scheme were created with Copilot, and the pipette-mixer image was drawn based from a photograph. (b) Fluorescence images captured 0, 24, and 48 h after dosing with 50 ng mRNA per well. Scale bar, 400 µm. (c–e) Green fluorescence intensity per cell, averaged across 6 images at each time point: (c) Comirnaty-like, (d) Spikevax-like, (e) Onpattro-like.

We next evaluated EGFP mRNA delivery mediated by these LNPs in EpiLiver, a polarized 3D primary human hepatocyte transwell model with distinct apical and basolateral surfaces and sustained liver-specific functions (Fig. 6, S7). LNPs were dosed basolaterally to better reflect hepatocyte exposure following systemic administration,^32^ and EGFP expression was monitored by live-cell fluorescence imaging. All 3 formulations produced detectable EGFP expression, with stronger EGFP signals at the higher mRNA dose and increasing expression over the observation period. Compared with HEK293T cultures, the 3D tissues showed a slower response and required more mRNA to produce detectable fluorescence. Together, these findings show that LNPs produced with the pipette-driven mixer support functional mRNA delivery in both HEK293T cells and a 3D human liver tissue model.

**Figure 6.**
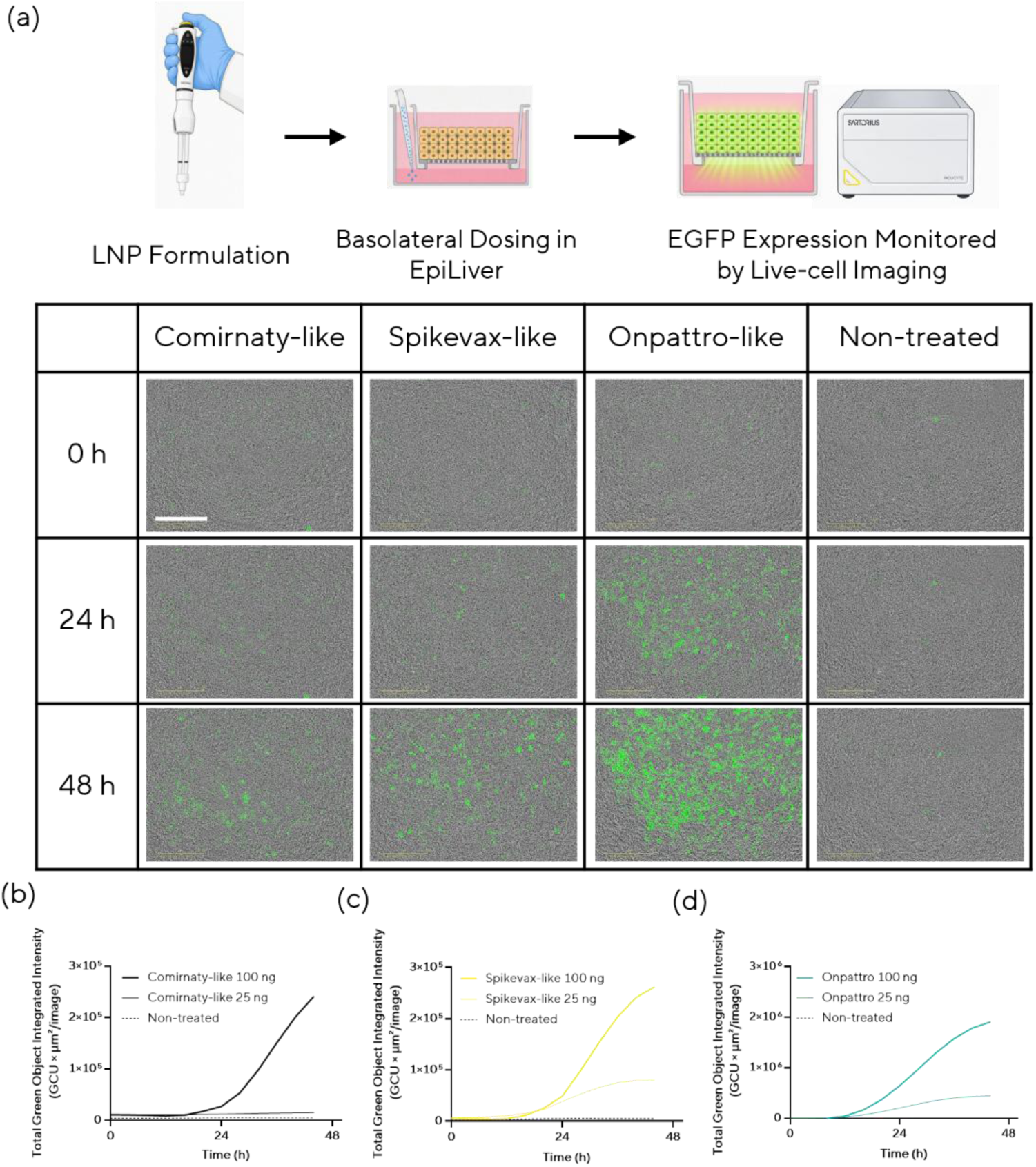
*In vitro* transfection of 3D liver tissue models with LNPs prepared using the pipette-driven mixer. LNPs were dosed basolaterally, and each sample was tested in four wells, with one image per well acquired every four hours. Illustrations in the scheme were created with Copilot; the pipette-mixer image was drawn from a photograph. (a) Fluorescence images acquired 0, 24 and 48 h after dosing (100 ng mRNA per well). Scale bar, 400 µm. (b–d) Total Green Object Integrated Intensity (GCU × µm²/image) in 3D cultures, quantified using the IncuCyte Basic Analyzer and averaged across 4 images at each time point over 48 h after dosing (25 and 100 ng mRNA per well): (b) Comirnaty-like, (c) Spikevax-like, (d) Onpattro-like.

To prepare formulations for the in vivo study, we next used firefly luciferase (Fluc) mRNA and evaluated formulation parameters in an automated workflow. The pipette-driven mixer was integrated with a liquid-handling robot capable of generating 96 LNP samples, each consisting of a 500 µL formulation followed by dilution with 3.5 mL PBS (Fig. 7a). The microfluidic mixer, dual-chamber pipette tip, and dilution tip were replaced after each preparation to minimize cross-contamination. We evaluated the effect of N/P ratio on particle size, PDI, and mRNA encapsulation across the three lipid formulations. The HTS study showed that low N/P ratios led to increased particle size and reduced encapsulation efficiency (Fig. 7b, c). An N/P ratio of 8 was selected for subsequent *in vivo* experiments based on the measured particle size and encapsulation efficiency.

**Figure 7.**
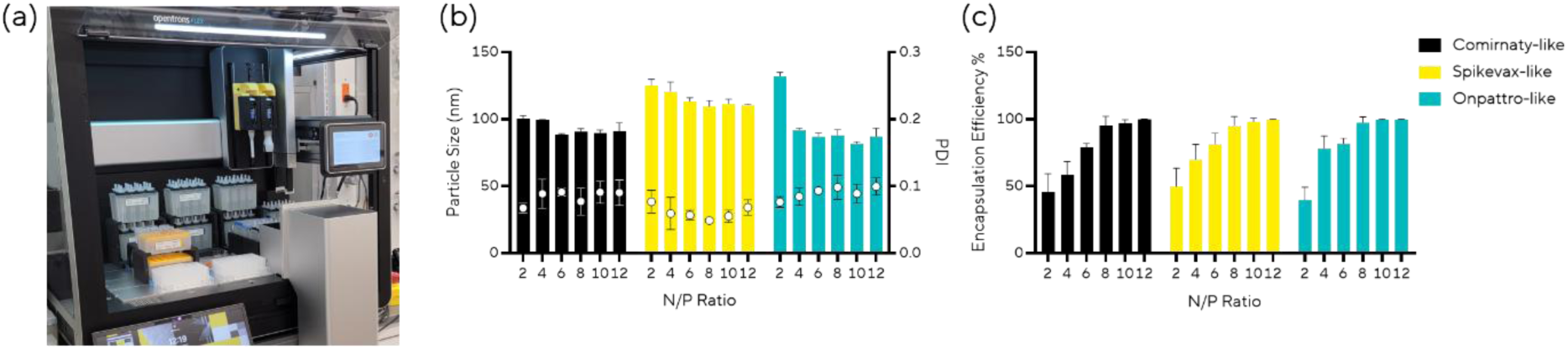
High-throughput screening (HTS) using a pipette mixer integrated into a liquid-handling robot. (a) Integration of the pipette mixer on the robotic platform. (b, c) HTS demonstration: the effect of the N/P ratio on mRNA encapsulation and particle size was assessed. Firefly luciferase (Fluc) mRNA was encapsulated across three LNP formulations. (b) Particle size and PDI. (c) mRNA encapsulation efficiency.

We next evaluated LNP-mediated mRNA delivery *in vivo* using Fluc mRNA encapsulated in three formulations, Comirnaty-like, Spikevax-like, and Onpattro-like. LNPs prepared using the high-shear pipette-driven mixer were administered intravenously to female C57BL/6 mice at doses of 2 or 10 μg mRNA per mouse (3 mice per group). Whole-body bioluminescence was measured at 1, 6, and 24 h after dosing (Fig. 8, S8). Luciferase expression was detected at 1 h, was highest at 6 h among the sampled time points, and declined by 24 h. Analysis of the 6 h data by two-way ANOVA revealed significant effects of formulation (p < 0.0001) and dose (p = 0.0004), as well as a significant formulation × dose interaction (p = 0.0009), indicating that the magnitude of the dose effect differed among formulations. Šídák-adjusted multiple comparisons detected higher bioluminescence at 10 µg than at 2 µg for the Comirnaty-like formulation (adjusted p < 0.0001 but not for the Spikevax-like or Onpattro-like formulations (adjusted p = 0.6142 and 0.9929, respectively). PBS-treated controls showed no detectable bioluminescence under the imaging conditions used. Ex vivo imaging of organs collected at 24 h showed that the liver was the predominant site of luciferase expression, with lower signals in the other examined organs. No overt signs of toxicity were observed at either tested dose during the 24 h observation period.

**Figure 8.**
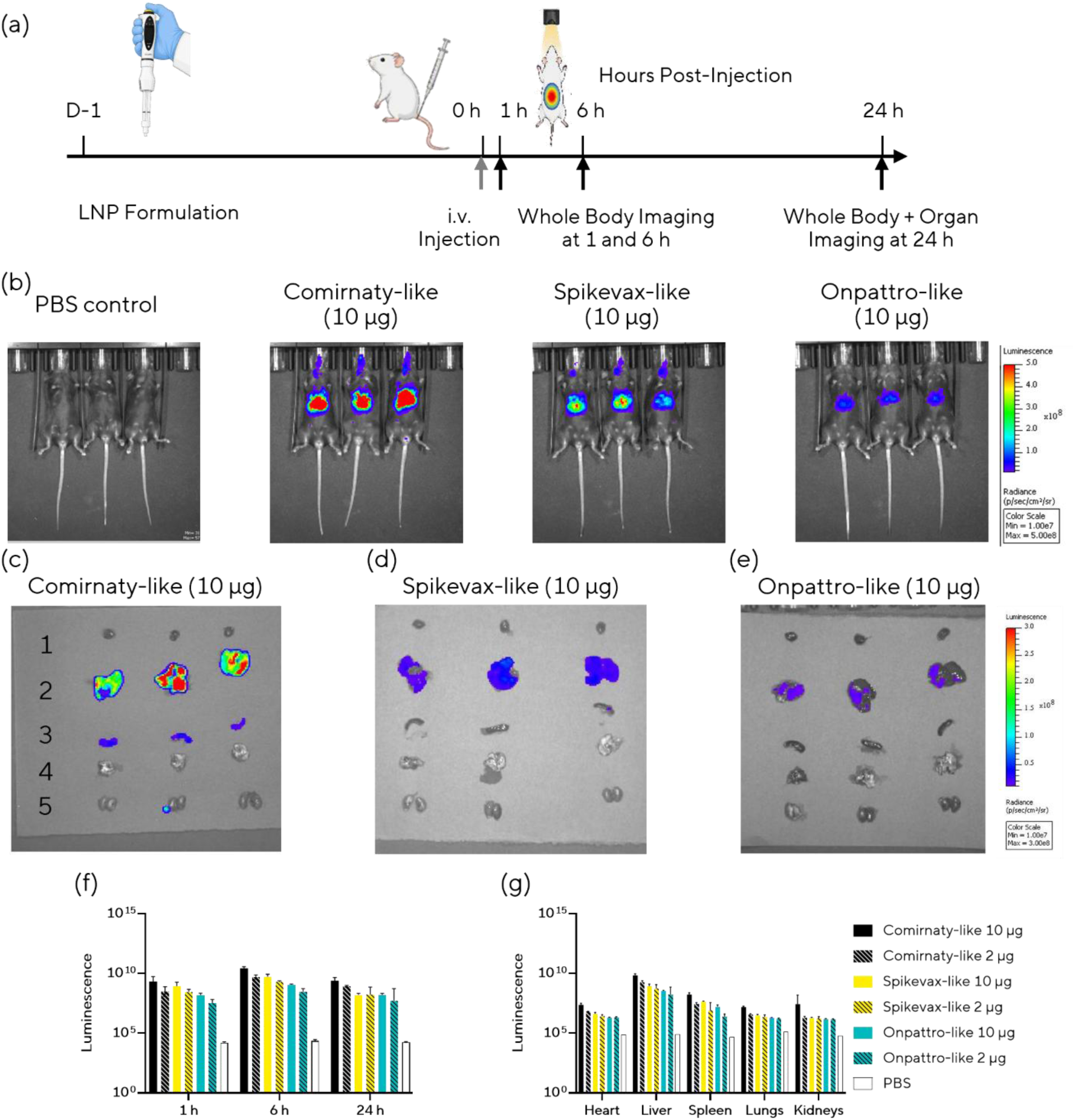
*In vivo* evaluation of LNP-mediated mRNA delivery. (a) Study design. Female C57BL/6 mice (n = 3) received intravenous injections of firefly luciferase mRNA–loaded LNPs prepared with the pipette mixer (2 and 10 µg mRNA per mouse); PBS-treated mice served as controls. Luciferase expression was monitored by IVIS imaging at 1, 6, and 24 h post-injection, and the heart, liver, spleen, lungs, and kidneys were collected at 24 h. (b) Representative IVIS images of LNP-treated mice at 6 h post-injection. (c–e) *Ex vivo* images of harvested organs: (1) heart, (2) liver, (3) spleen, (4) lungs, and (5) kidneys. (f) Time course of luminescence. (g) Luminescence in harvested organs. All data are presented as mean ± standard deviation.

### Mixer Scale-Up and Validation of Large-Scale LNP Production

Scaling microfluidic LNP production remains challenging because increasing throughput or changing mixing platforms can alter mixing conditions and, consequently, LNP properties and biological performance.^9,33^ To address this challenge, we investigated whether scaling up the pipette mixer geometry could increase production throughput while maintaining comparable particle size and mRNA encapsulation. For this evaluation, we developed a large-scale LNP production system with integrated in-line dilution using an aqueous buffer (Fig. 9a). To increase volumetric throughput, we selected a four-fold geometric scale-up that balanced mixing requirements with back pressure and pumping-system capabilities. The scaled geometry was designed to operate below 3 bar within a flow-rate range compatible with the pumping system.

**Figure 9.**
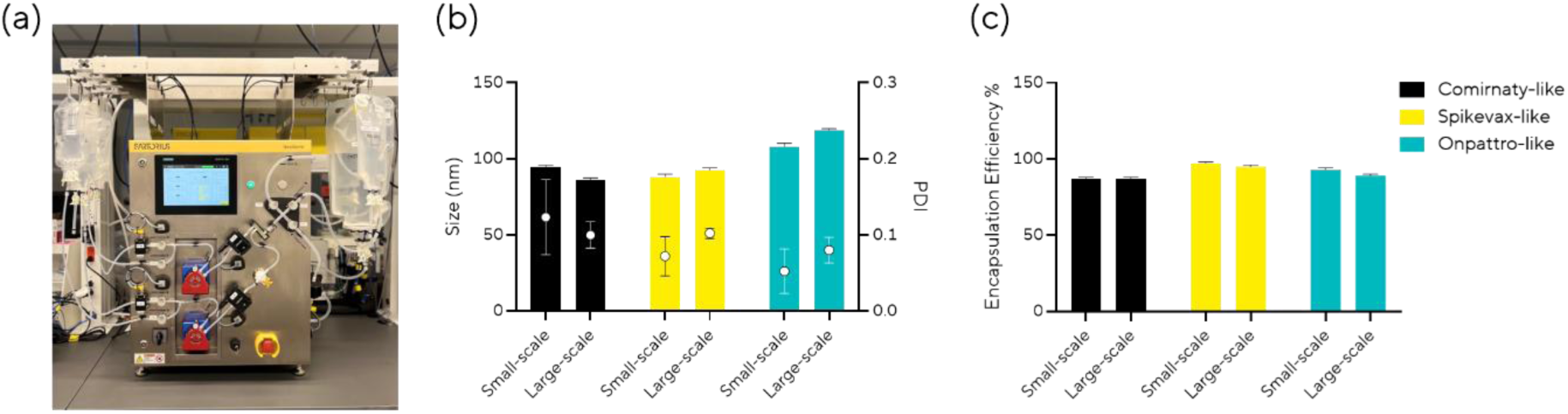
Scale-up production of LNPs. (a) The large-scale LNP production unit. (b, c) Demonstration of large-scale LNP production with similar particle characteristics to one with small-scale pipette-driven mixer: (b) particle size and (c) mRNA encapsulation.

To identify operating conditions expected to preserve a characteristic mixing time comparable to that of the small-scale device, we applied a dimensionless scaling analysis. The characteristic mixing time was defined as the time required for convective stretching to thin concentration filaments to the diffusive length scale, where molecular diffusion can complete homogenization. The characteristic mixing time, derived in the Supporting Information, is given by:

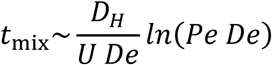

For the scaled-up geometry, the hydraulic diameter was increased four-fold: *D*′*_H_* = 4*D_H_*

Accordingly, the dimensionless groups scale as

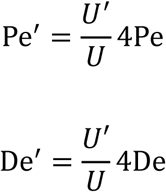

where Pe is the Peclet number, U is the average velocity, and De is the Dean number. Primed quantities denote the corresponding values for the large-scale geometry. This framework was then used to identify the flow rate in the large-scale device that would preserve a mixing time comparable to that of the small-scale geometry.

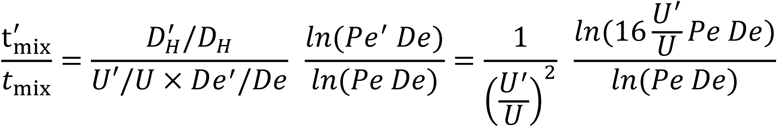

For Pe = 300000, De = 150, and 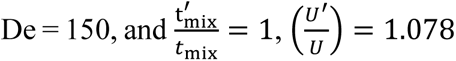. The flow rate in the scale-up chip is given by *Q*^′^ = *A*^′^*U*^′^ = 16*A* × 1.078 *U* = 17.2 *Q*. Therefore, achieving comparable mixing in the scaled-up chip requires a flow rate approximately 17 times higher than that used in the small-scale chip. Consistent with this estimate, the following study produced LNPs at 150 mL/min, which is 16.6-fold higher than the average flow rate used for small-scale production (9 mL/min). CFD was used to calculate the mixing index, a metric of global mixing performance, as well as temporal changes in velocity and concentration in the small- and large-scale geometries. These analyses were performed to evaluate overall homogeneity and start-up transients, as shown in Fig. S9. Because the velocity profiles were similar between the two chip scales and the chip dimensions were increased four-fold, the large-scale geometry required approximately four times longer to reach a steady-state concentration field (Fig. S9c). However, as discussed in the first section and the following paragraph, micromixing time was more directly relevant to LNP quality than the global mixing metrics evaluated here.

Using our scale-up equipment, we produced 75 mL of three benchmark LNPs (Comirnaty-like, Spikevax-like, and Onpattro-like). Both small- and large-scale preparations were dialyzed after formulation, but only the large-scale preparations underwent in-line dilution before dialysis. The large-scale preparations exhibited particle size, PDI, and mRNA encapsulation efficiency similar to those obtained with the small-scale pipette-driven mixer, supporting scale-up with respect to these measured attributes (Fig. 9b, c).

## Conclusions

In this study, we introduced a single-use, disposable, pipette-driven microfluidic mixer that addresses key practical barriers in LNP formulation development, including equipment complexity, limited automation compatibility, material consumption, and scale-up uncertainty. The platform reproducibly produced LNPs that achieved functional mRNA delivery in both *in vitro* and *in vivo* models, while integration with a liquid-handling robot enabled high-throughput formulation screening using minimal material with low batch-to-batch variability. The mixer geometry was also scalable, enabling larger-scale production of LNPs with particle size, PDI, and encapsulation efficiency comparable to those obtained at small scale. Together with orthogonal validation of particle quality, these findings establish the pipette-driven mixer as an accessible, scalable, and automation-compatible platform for bridging early formulation screening and larger-scale LNP production with reduced process development effort.

## Experimental Section

### Materials

The plasmid DNA encoding EGFP used was supplied by Aldevron and subsequently linearized using BspQI (Hongene). EGFP mRNA was synthesized from the linearized DNA in-house, with *in vitro* transcription reagents sourced from Hongene. The IVT protocol was adapted from Boman *et al*.^34^ The synthesized mRNA was purified using the Monarch Spin RNA Cleanup Kit (New England Biolabs) for small-scale preparations and an oligo dT column (Sartorius) with an AKTA chromatography system (Cytiva) for large-scale chromatography-based purification. The concentration and purity of the mRNA were assessed using a Nanodrop One spectrophotometer (Thermo Fisher).

SM-102 and D-Lin-MC3-DMA (MC3) lipids were purchased from Ambeed; ALC-0315 from BroadPharm; DMG-PEG2000 and ALC-0159 from Avanti Research; 1,2-distearoyl-sn-glycero-3-phosphocholine (DSPC) from Lipoid; and cholesterol from Sigma.

### Mixer equipment

#### Microfluidic Pipette Mixer

The microfluidic pipette mixer comprised a Picus® 2 electronic positive-displacement pipette (Sartorius Liquid Handling Oy) with adjustable dispensing speed, a single-use dual-chamber pipette tip, and a single-use microfluidic LNP mixer fabricated using a Profluidics 285D printer (CADworks3D). The dual-chamber pipette tip was attached to the Picus® 2 pipette, and lipid and mRNA solutions were transferred to separate wells of a 24-well plate for aspiration. The two outlets of the dual-chamber tip were then inserted into the inlets of the microfluidic LNP mixer. The solutions were dispensed through the mixer to form LNPs, which were collected from the outlet in a 1.5 mL microcentrifuge tube. The system produces 500–2500 µL per run and is compatible with both manual and automated workflows.

#### Large-scale LNP Production Equipment

LNP production was scaled up using a prototype large-scale LNP mixer equipped with a single-use flow kit and a mixing chamber that matches the geometry and flow profile of the pipette mixer. Lipid and mRNA solutions, as well as PBS and ethanol, were filled into Flexsafe storage bags (Sartorius). The system enables continuous production of 25 mL to 10 L of LNPs at flow rates of 40–400 mL/min, with in-line dilution using PBS. The resulting LNPs were collected in a Flexsafe bag.

#### Integration of the Pipette Mixer with a Liquid-handling Robot

An Opentrons Flex was configured with a Picus® 2 electronic positive-displacement pipette for LNP production and a single-channel Picus® 2 (1 mL) pipette for post-synthesis dilution. The Flex deck (15 available slots, with one of the 16 slots occupied by the waste chute) accommodated disposable dual-syringe assemblies, single-use mixer chips, pipette tips, 48-well plates for lipid and mRNA solutions, 48-deep-well plates for sample collection, and 50 mL tubes for PBS. Each sample was prepared as described in the Microfluidic Pipette Mixer section and collected in a 48-deep-well plate, with two collection plates accommodating up to 96 samples. The single-channel Picus® 2 then added 3.5 mL PBS to each 500 µL LNP preparation. To minimize cross-contamination, dual-syringe assemblies, mixer chips, and pipette tips were replaced for every preparation.

#### Computational Fluid Dynamics (CFD) Analysis

Computational fluid dynamics (CFD) simulations were performed using MStar-CFD to quantify the mixing efficiency of each system. MStar-CFD performed direct numerical simulations based on the lattice Boltzmann Method (LBM), which has been validated in several publications.^35,36^ A fixed time step of 3×10^-7^ s was used. The fluid was modeled as Newtonian, with viscosity 1 cP and density 1000 kg m^-3^. A second component was modeled as a passive scalar introduced at a molar flow rate of 1 mol s^-1^ with a constant diffusion coefficient value of 1×10^-9^ m^2^ s^-1^.

The microchannel was operated at a volumetric flow rate of 1.5 × 10⁻⁷ m³ s⁻¹. The channel height and width were 2.0 × 10⁻⁴ m and 8.1 × 10⁻⁴ m, respectively, corresponding to a cross-sectional area of 1.62 × 10⁻⁷ m² and a hydraulic diameter (*D_H_*) of 3.2 × 10⁻⁴ m, and an average flow velocity of 0.926 m s⁻¹. For the small-scale geometry, a lattice resolution of L_y_ = 30 cells was used, corresponding to a lattice spacing of Δx = 6.67×10^-6^ m. The computational domain contained approximately 72.8 million lattice cells. For the large-scale geometry, a lattice resolution of L_y_ = 45 cells across the channel height was applied, corresponding to a lattice spacing of Δx = 1.78×10^-5^ m. The computational domain contained approximately 261.1 million lattice cells. Both simulations were performed at a Courant number ≤ 0.01 and run for 1 s of physical time, which was sufficient for transient effects to dissipate and steady-state flow conditions to be established.

The Peclet number is calculated as 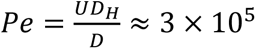, indicating strongly advection-dominated transport. Assuming working fluid with a kinematic viscosity of approximately 10^-6^ m^2^ s^-1^, the Reynolds number is estimated as Re ≈ 300. Under these conditions, the flow remained laminar while exhibiting non-negligible inertial effects. Because of the curved geometry, secondary Dean vortices were expected to enhance transverse transport and mixing. The influence of secondary flow structures was evaluated using the Dean number, which was approximately 150. This relatively high Dean number indicates the presence of strong secondary flow that can substantially increase cross-sectional mixing and reduce diffusion-limited transport effects.

Resolved shear was derived from local velocity gradients obtained from the LBM solution of the Navier–Stokes equations. At each lattice cell, the local strain-rate tensor was computed from the macroscopic velocity field, obtained as moments of the particle distribution functions, as 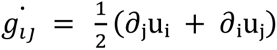. For a Newtonian fluid, the shear stress was calculated as 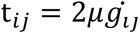. The scalar mean resolved shear value reported in post-processing is the volume- or area-averaged magnitude of the shear stress:

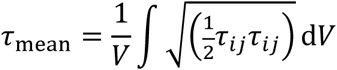

This formulation quantifies the intensity of local velocity gradients and provides a measure of hydrodynamic stress distribution within the simulated domain.

The mixing index was calculated from the concentration standard deviation using the following equation:

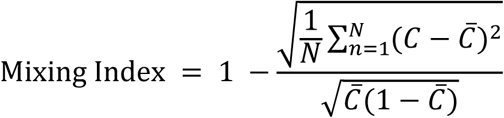

where *C_i_* is the concentration of the *i*-th cell and *C̅* is the mean concentration.

Micromixing time distribution was calculated based on a correlation fitted to Kolmogorov theory developed by Nouri *et al*:

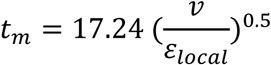

where *v* is the kinematic viscosity and *ε_local_* is the energy dissipation rate at any given computation cell.^37^

#### Preparation of Lipid Nanoparticles (LNPs)

Lipids were dissolved in ethanol, and the aqueous phase consisted of a 10 mM citrate buffer at pH 4 (Teknova). The flow rate ratio was set to 3, the N/P ratio to 6, and the initial lipid concentration to 10 mM. Three LNP formulations tested were summarized in Table 1. Samples were prepared using the pipette-driven mixer or 10 cycles of manual pipetting, then dialyzed overnight against PBS (Boston BioProducts) in Slide-A-Lyzer MINI dialysis devices (Thermo Fisher) with a 10 kDa molecular weight cutoff. The LNP samples prepared using the HTS equipment were analyzed after dilution in PBS.

### Characterization of LNPs

#### Particle Size and mRNA Loading

Particle size was measured using a Zetasizer Pro (Malvern Panalytical), and mRNA loading was quantified using RiboGreen assay (Thermo Fisher). Encapsulation efficiency (*EE*) was calculated using the formula: *EE* = (1 − *C_f_*⁄*C_t_*) × 100 (%), where *C_f_* represents the free mRNA concentration and *C_t_* the total mRNA concentration. Total mRNA concentration was measured by disrupting the lipid membrane with Triton X-100 (Sigma), while the free mRNA concentration was measured without addition of Triton X-100.

#### Nanoparticle Tracking Analysis (NTA)

Particle concentration of LNPs was calculated using NS300 NanoSight (Malvern Panalytical) as reported before.^24^ NTA measurements used a 532 nm, 45 mW green laser with an sCMOS camera. Five 30-second videos were recorded per sample at camera level 16 and screen gain 1 (software v3.4), under these conditions: cell temperature 25°C, syringe speed 35. Videos were analyzed with a detection threshold of 5 and screen gain 10; advanced settings were set to auto. Each sample was tested in triplicate, with PBS as the dilution buffer. After establishing the optimal scattering dilution (at least 7 × 10⁷ particles/ml), a concentration double that amount was used to prepare triplicates for fluorescence mode. Fluorescent-based NTA was performed by labeling LNPs using CellMask Orange (Thermo Fisher) before the NTA measurement. Each sample was incubated with 5 µg/ml of dye for 30 minutes in the dark at room temperature. The triplicates were analyzed using the same parameters established for scattering mode.

#### Asymmetric-Flow Field-Flow Fractionation (AF4-MALS/QELS)

Asymmetric flow field-fractionation (AF4) measurements were performed using an Eclipse AF4 system (Wyatt Technology) equipped with a dilution control module, a long, variable-height channel with a 350 μm spacer, and a regenerated cellulose membrane with a molecular weight cutoff of 10 kDa (Wyatt Technology).^26^ The system was coupled to a DAWN multi-angle light scattering (MALS) detector equipped with an integrated WyattQELS dynamic light scattering (DLS) module, an Optilab differential refractive index (dRI) detector (Wyatt Technology), and a Waters 2998 photodiode array (PDA) detector (Waters Corporation) monitoring UV absorbance at 260 and 280 nm. The channel temperature was set at 25 °C using a channel thermostat. The autosampler temperature was set to 4°C. Prior to the sample analysis, the membrane was conditioned by injecting 50 µL of bovine albumin serum (BSA) at 2 mg/mL. The angular responses of the MALS detectors were normalized relative to the 90° scattering angle using monomeric BSA as an approximately isotropic scattering standard, according to the manufacturer’s recommendations. Undiluted (neat) samples were analyzed in triplicate using an injection volume of 30 μL.

The mobile phase consisted of 1× phosphate-buffered saline (PBS, pH 7.4). The channel flow, detector flow, and injection flow rates were set to 1.6, 0.5, and 0.2 mL/min, respectively, with the focus position maintained at 25%. The AF4 separation program consisted of an initial elution step at zero crossflow (0-2 min), followed by focusing at a constant crossflow rate of 1.5 mL/min (2-4 min), and focusing with sample injection at the same crossflow rate (4-7 min). Subsequently, the cross flow was decreased exponentially from 1.5 to 1.0 mL/min (7-17 min) and from 1.0 to 0.05 mL/min (17-37 min), followed by a linear decrease from 0.05 to 0 mL/min (37-39 min). The program concluded with a final 3 min step at zero crossflow (39-42 min).

Data analysis was performed using ASTRA software (version 8.3.0, Wyatt Technology). The hydrodynamic radius (R_h_) was determined from online DLS measurements using the QyattQELS module, while the radius of gyration (R_g_) was calculated from MALS data using a Berry fit. The R_g_/R_h_ ratio was calculated as a shape factor to assess particle shape and morphology. For each injection, mean values of R_g_, R_h_, and R_g_/R_h_ were calculated from data points within the full width at half maximum (FWHM) of the elution peak. The FWHM was determined from the 90° MALS scattering intensity profile using the retention times corresponding to 50% of the maximum peak height. Data points outside the FWHM were excluded to minimize contributions of low-signal noise, particularly in the online DLS measurements. The resulting mean values obtained from three replicate injections were subsequently averaged, and standard deviations were calculated across the replicate injections.

#### Single Particle Automated Raman Trapping Analysis (SPARTA)

LNP lipid composition was analyzed using a SPARTA AGIS I benchtop instrument (SPARTA Biodiscovery).^24,27^ Instrument control and data acquisition were performed in SPARTA Control v 1.1.3. LNP samples were dialyzed overnight against PBS, further diluted 60-fold in PBS before analysis. A 120 μL aliquot of particle suspension (1 × 10^10^–1 × 10^12^ particles mL^-1^) was dispensed onto a SPARTA Sample Slide which was then loaded into the AGIS I system. All measurements were carried out at ambient temperature (20 ± 1 °C). Individual particles were held for 10 s per trap whilst their Raman signal was acquired after which the laser shutter was closed for 1 s to release the trapped particle and allow diffusion of a new particle into the confocal volume. Blank 2 mM tris(hydroxymethyl)aminomethane) (Thermo-Fisher Scientific) and sodium chloride (Thermo-Fisher Scientific) was measured for 20 acquisitions at 10 s and the average was used for background subtraction.

Sample data of approximately 250 particles was analyzed using in-house SPARTA Discovery software v.1.1.3. A spectral response correction was applied, cosmic spikes were removed using Z limit 20, empty particle trapping events were removed through thresholding, and a background subtraction of 95% was applied. All spectra were then truncated to a ROI of 416-1800 cm^-1^, followed by a baseline correction with smoothness factor 7 and differential order 2. The data was then smoothed using a filter with polynomial order 2 and window 7 before being normalized by setting the area under each spectrum to 1.

#### Multicolor CICS Analysis of LNP Composition and mRNA Payload

Single-particle analysis of lipid nanoparticles (LNPs) was performed using a three-color cylindrical illumination confocal spectroscopy (CICS) platform based on the previously described design.^29–31^ Briefly, fluorescence excitation was provided by fiber-coupled lasers operating at 488, 561, and 638 nm (Cobolt Skyra, HÜBNER Photonics). A cylindrical lens shaped the excitation beam into a one-dimensional light sheet, which was focused into a fused-silica microcapillary (4 µm inner diameter, Polymicro) using a 100× oil-immersion objective (UPLFLN, numerical aperture=1.3, Olympus). Fluorescence emission was collected through the same objective, passed through a quad-bandpass filter and a confocal pinhole, and directed to single-photon avalanche photodiode detectors (SPAD, SPCM-AQRH10, Excelitas), with one detector for each fluorescence channel. The fluorescence detection bands were 525/28, 575/25, and 676/37 nm, expressed as center wavelength/bandwidth.

The mRNA encoding green fluorescent protein (EGFP) (ApexBio) was covalently labeled with cyanine 5 (Cy5). To label the lipid component, TopFluor-labeled phosphatidylcholine (TopFluor-PC; Avanti Polar Lipids) was incorporated into the LNP formulations by replacing a portion of the unlabeled DSPC helper lipid. The TopFluor-PC content was 0.5%.

Individual fluorescence bursts were characterized by their event time, peak intensity, width (*i.e.*, duration, millisecond), and size (*i.e.*, total photon count). Temporal coincidence between the TopFluor and Cy5 fluorescence channels was used to classify detected events as mRNA-loaded LNPs (TopFluor⁺/Cy5⁺), empty LNPs with no detectable mRNA signal (TopFluor⁺/Cy5⁻), or free mRNA (TopFluor⁻/Cy5⁺). The empty-LNP fraction was calculated as the number of empty LNPs divided by the total number of detected LNPs, including both empty and mRNA-loaded LNPs.

The mRNA payload distribution was estimated by fluorescence-distribution deconvolution using the previously described approach.^29,38^ The experimentally measured Cy5 fluorescence distribution of individual mRNA molecules served as the single-mRNA reference. Basis distributions representing LNPs containing different integer numbers of mRNA molecules were generated by scaling the fluorescence values of this reference distribution by the corresponding copy number. The measured Cy5 fluorescence distribution of mRNA-loaded LNPs was then fitted as a nonnegative weighted sum of these basis distributions. The normalized fitted weights yielded the estimated fraction of mRNA-loaded LNPs containing each number of mRNA molecules.

### Cryogenic electron microscopy (Cryo-EM)

LNPs were visualized by cryo-EM as reported before.^24^ LNP samples were dialyzed overnight in PBS and then diluted for imaging. Each sample was preserved in vitrified ice supported by holey carbon films on 400-mesh copper grids. Each sample was prepared by applying a drop of sample suspension to a cleaned grid, blotted away with filter paper, and immediately followed with vitrification in liquid ethane. Grids were stored under liquid nitrogen until transferred to the electron microscope for imaging. Micrographs were collected on a Glacios Cryo (Thermo Fisher) at 200 kV with a Falcon 4 direct electron detector. The images were acquired at a nominal underfocus of −7.0 to −4.5 μm and electron doses of 2–12 e^-^/Å^2^.

### *In Vitro* Transfection Assay

*In vitro* transfection activity was evaluated as previously described.^39^ For these studies, LNPs encapsulating CleanCap EGFP mRNA (TriLink) were prepared using the high-shear mixer. HEK293T cells (ATCC) were cultured in DMEM (Cytiva) supplemented with 10% fetal bovine serum (Gibco), 1% GlutaMax (Gibco), and 1% penicillin–streptomycin (Gibco), and seeded in 96-well plates 24 h before dosing. Time-lapse fluorescence images were acquired hourly for 72 h after dosing using an IncuCyte S3 (Sartorius). Each sample was tested in three wells, with two images acquired per well at each time point. Cell viability and transfection, measured as green fluorescence intensity in live cells, were analyzed using the IncuCyte AI Cell Health Analysis Software Module version 2025B.

A 3D liver tissue model (EpiLiver, MatTek) was also used to assess *in vitro* transfection.^32^ LNP samples were applied to the basolateral side, and transfection was monitored over a 48-hour culture period on an IncuCyte SX5 by measuring green fluorescence intensity throughout the culture period. Each sample was tested in four wells, with one image acquired per well every four hours. Analysis was performed with Basic Analyzer version 2025C with Beta version Mattek Vessel Definition.

### *In Vivo* Transfection Assay

Firefly luciferase (Fluc) mRNA (Hongene) was encapsulated in LNP formulations modeled after Comirnaty, Spikevax, and Onpattro. LNPs were prepared with the high-shear mixer, dialyzed against PBS, concentrated using Vivaspin concentrators (Sartorius), and sterile-filtered. Six- to seven-week-old female C57BL/6 mice (n = 3 per group) received a single intravenous dose of 10 or 2 µg mRNA per animal. Whole-body transfection was assessed by IVIS imaging at 1, 6, and 24 h post-dose, and the heart, liver, lungs, kidneys, and spleen were harvested for *ex vivo* imaging at 24 h post-dose.

### Data Analysis

Data analysis was performed in GraphPad Prism (version 11).

## Acknowledgement

We acknowledge Sartorius Liquid Handling team for providing us with the custom-made Picus 2 pipettors. The authors appreciate insightful discussions and resources from T. Kwon, A. Larey, and the Sartorius Corporate Research group. We thank J. Sun, A. Li, G. Datto, J. Wang, J. Ma, D. Yu, and K. Hsieh at Johns Hopkins University for technical support with CICS and *in vivo* assays. We are grateful to J. Penders, A. Leach, and the SPARTA Biodiscovery team for their technical support and valuable insights. The authors acknowledge C. Calloway for electron microscopy, which was performed in the Electron Microscopy Resource (RRID: SCR_012366) in the Center for Advanced Research Technologies at the University of Rochester Medical Center.

## Supporting Information Available

Video demonstration of pipette-driven mixer operation; CFD micromixing statistics; HPLC analysis of mRNA purification; NTA, AF4–MALS/QELS, CICS, SPARTA, cryo-EM, *in vitro* transfection, and *in vivo* imaging data; dimensionless mixing-time analysis; scale-up CFD comparison; supplementary references.

## Notes

Takeda, F. Wang, Gu, Forster, Hou, Hemmati, Partopour, Finelli, Molaei, Shafiee, Maugeri, Kim, Apel, Norton, Chiang, Mayo, Ayehunie, Akbari, and Dehghani were employees of Sartorius companies during this study; eight authors are no longer affiliated with Sartorius companies at the time of submission; they received salary support from Sartorius companies while the work was conducted. Patent applications related to this work have been filed. Sartorius has a commercial partnership with, and is an investor in, SPARTA Biodiscovery. The authors declare no additional competing interests.

## Author Contributions

Takeda: Investigation (lead), Methodology (supporting), Resource (supporting), Writing/ original draft (lead), review and editing (lead), Visualization (lead), Supervision (supporting).

F. Wang: Investigation (supporting), Methodology (supporting), Resource (lead), Writing/ review and editing (supporting).

Gu: Investigation (supporting), Methodology (supporting), Writing/ review and editing (supporting).

Forster: Investigation (supporting), Methodology (supporting), Writing/review and editing (supporting).

Hou: Investigation (supporting), Methodology (supporting), Writing/review and editing (supporting).

Hemmati: Investigation (supporting), Methodology (supporting), Visualization (supporting), Writing/ original draft, review and editing (supporting)

Partopour: Investigation (supporting), Methodology (supporting), Visualization (supporting), Writing/ original draft, review and editing (supporting)

Finelli: Investigation (supporting), Methodology (supporting), Writing/review and editing (supporting).

Molaei: Resource (supporting), Methodology (supporting), Writing/ review and editing (supporting).

Shafiee: Resource (supporting), Writing/ review and editing (supporting). Maugeri: Investigation (supporting), Writing/review and editing (supporting).

Kim: Investigation (supporting), Methodology (supporting), Writing/review and editing (supporting).

Apel: Investigation (supporting), Methodology (supporting), Writing/ original draft, review and editing (supporting).

Norton: Investigation (supporting), Methodology (supporting), Writing/review and editing (supporting).

Li: Resource (supporting), Investigation (supporting), Methodology (supporting), Visualization (supporting), Writing/ original draft (supporting), review and editing (supporting), Supervision (supporting).

Lin: Resource (supporting), Investigation (supporting), Methodology (supporting), Visualization (supporting), Writing/ original draft (supporting), Writing/review and editing (supporting).

Chiang: Investigation (supporting), Methodology (supporting), Visualization (supporting).

Mayo: Resource (supporting), Methodology (supporting), Supervision (supporting). Writing/review and editing (supporting).

Ayehunie: Methodology (supporting), Funding Acquisition (supporting), Supervision (supporting). Writing/review and editing (supporting).

Akbari: Conceptualization (equal), Funding Acquisition (equal), Writing/review and editing (supporting), Supervision (supporting).

Mao: Conceptualization (supporting), Resource (supporting), Writing/review and editing (supporting), Supervision (supporting)

T.H. Wang: Resource (supporting), Writing/review and editing (supporting), Supervision (supporting)

Dehghani: Conceptualization (equal), Funding Acquisition (equal), Methodology (lead), Visualization (supporting), Writing/original draft, review and editing (supporting), Supervision (lead)

## Data Availability Statement

The data that support the findings of this study are available within the article and from the corresponding author upon reasonable request.

