## Supporting Information for "An Accessible and Scalable Pipette-Driven Microfluidic Platform for Lipid Nanoparticle Formulation"

Table of Contents


| | |
|---|---|
| Figure S1. Video demonstrating the operation of the pipette-driven microfluidic mixer | 3 |
| Table S1. Summary statistics for nonzero micromixing time values calculated by CFD | 4 |
| Figure S2. PATfix analytical chromatography analysis confirming the removal of unincorporated nucleotides from in-house-synthesized mRNA after purification. | 4 |
| Figure S3. Particle size distribution of Comirnaty-like, Spikevax-like, and Onpattro-like formulations measured using scattering- and fluorescence-based NTA | 5 |
| Figure S4. Asymmetric field-flow fractionation analysis showing shape factor and MALS signal as functions of elution time for LNPs prepared using four mixer geometries | 6 |
| Figure S5. Scatter plots obtained by CICS analysis | 7 |
| Table S2–S10. SPARTA analysis of particle-level Raman shift distributions. | 8 |
| Table S11–S13. mRNA payload distribution per particle analyzed by CICS. | 11 |
| Figure S6. *In vitro* transfection in HEK293T cells | 12 |
| Figure S7. *In vitro* transfection of 3D liver tissue models with LNPs prepared by the high-shear mixer | 13 |
| Figure S8. Representative IVIS images of female mice treated with LNPs at each time point. | 14 |
| Dimensionless analysis of the mixing time | 14 |
| Figure S9. CFD simulation comparing mixing performance between the scale-up mixer and small-scale mixer | 17 |
| References | 17 |

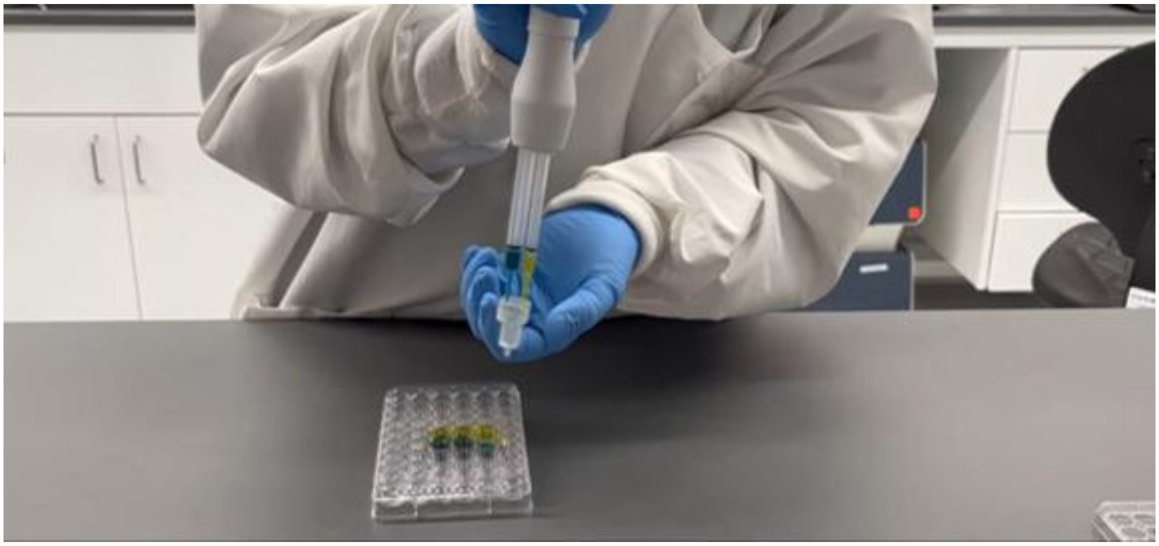

Figure S1. Video demonstrating the operation of the pipette-driven microfluidic mixer.

Multimedia available online.

Table S1. Summary statistics for nonzero micromixing time values calculated by CFD. SD, standard deviation. CQV, coefficient of quartile variation.

| Mixer | Non-zero (%) | Mean (s) | SD (s) | Median (s) | CQV |
|---|---|---|---|---|---|
| **Low-Shear** | 35.1% | 0.0715 | 1.27 | 0.0322 | 0.63 |
| **High-Shear** | 29.1% | 0.00254 | 0.00209 | 0.00212 | 0.21 |

mRNA Analysis by Analytical Chromatography

The synthesized mRNA was analyzed using PATfix LNP Switcher Module and IVT Monitoring Modules (Sartorius).[1,2] The mRNA was characterized before and after purification process using spin columns (New England Biolabs) and chromatography.

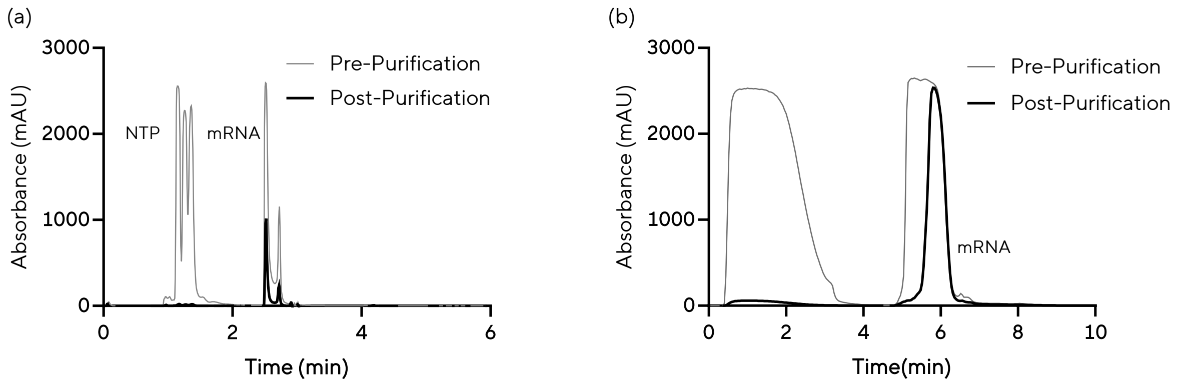


Figure S2. PATfix analytical chromatography analysis confirming the removal of unincorporated nucleotides from in-house-synthesized mRNA after purification. (a) The IVT monitoring module. (b,c) LNP Switcher module.

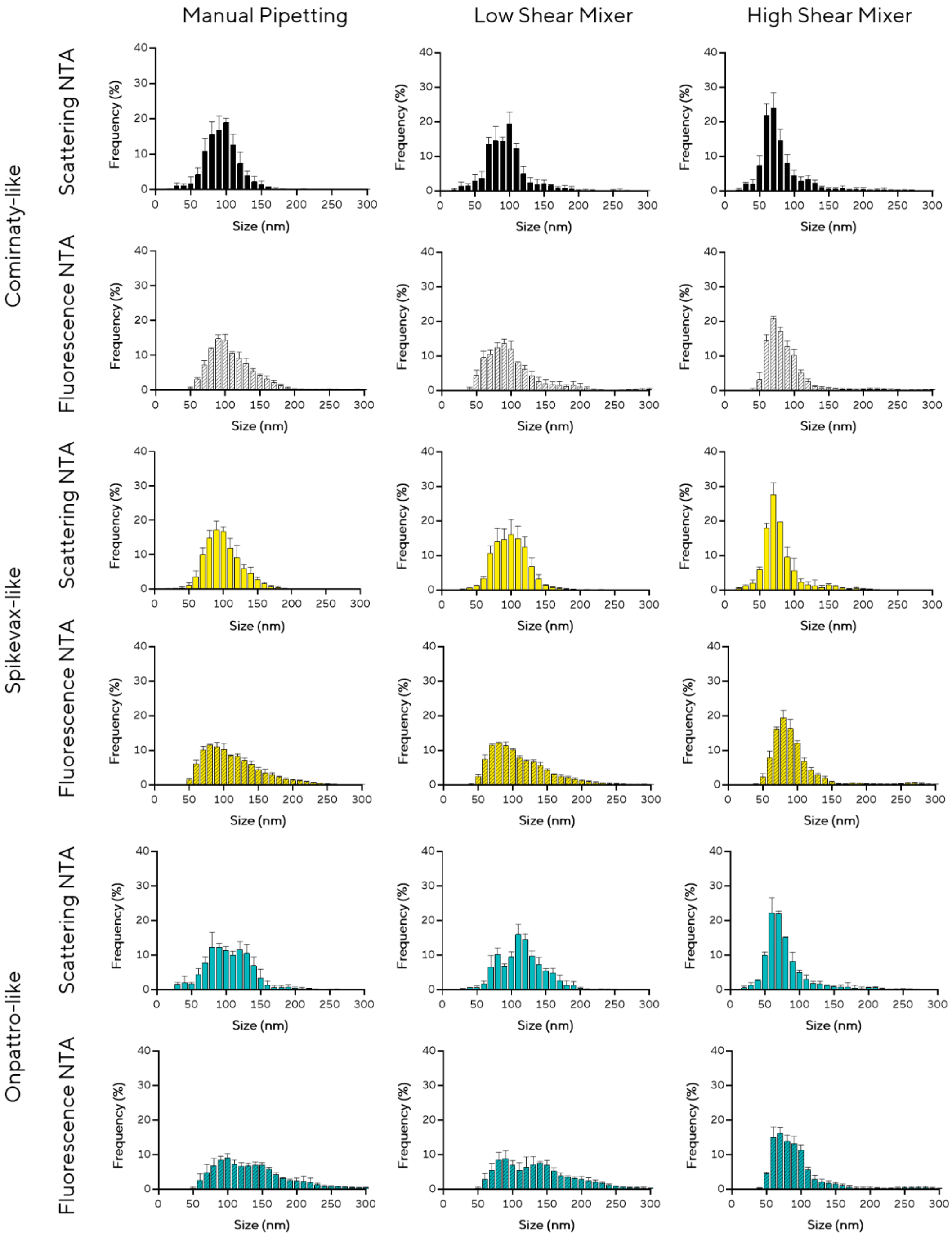


Figure S3. Particle size distribution of Comirnaty-like, Spikevax-like, and Onpattro-like formulations, measured using scattering- and fluorescence-based NTA.

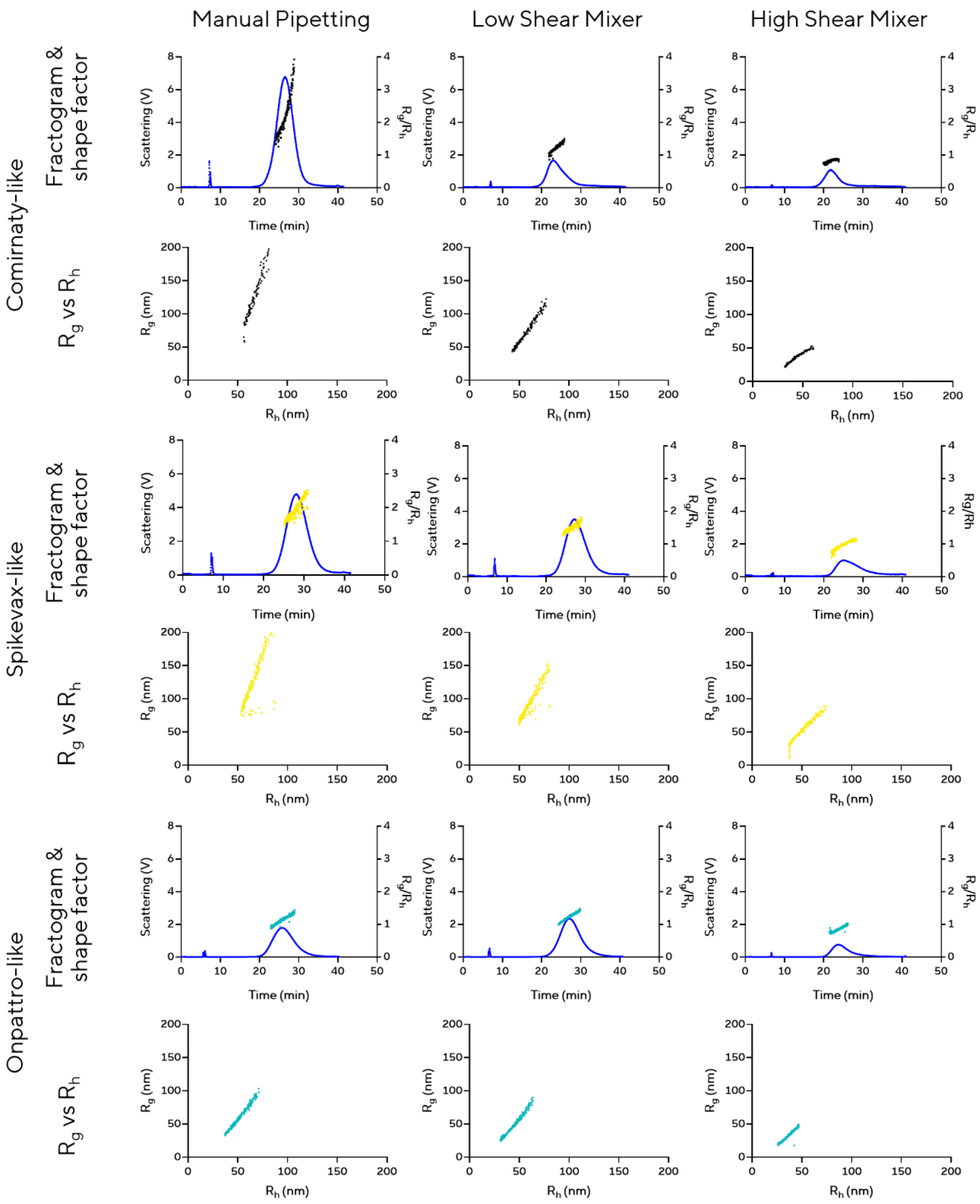


Figure S4. Asymmetric field-flow fractionation (AF4) analysis of LNPs prepared under three mixer conditions. The upper panels show the shape factor (right axis, $R_g/R_h$) and the MALS scattering signal (left axis) as functions of elution time. The lower panels show $R_g$ against $R_h$.

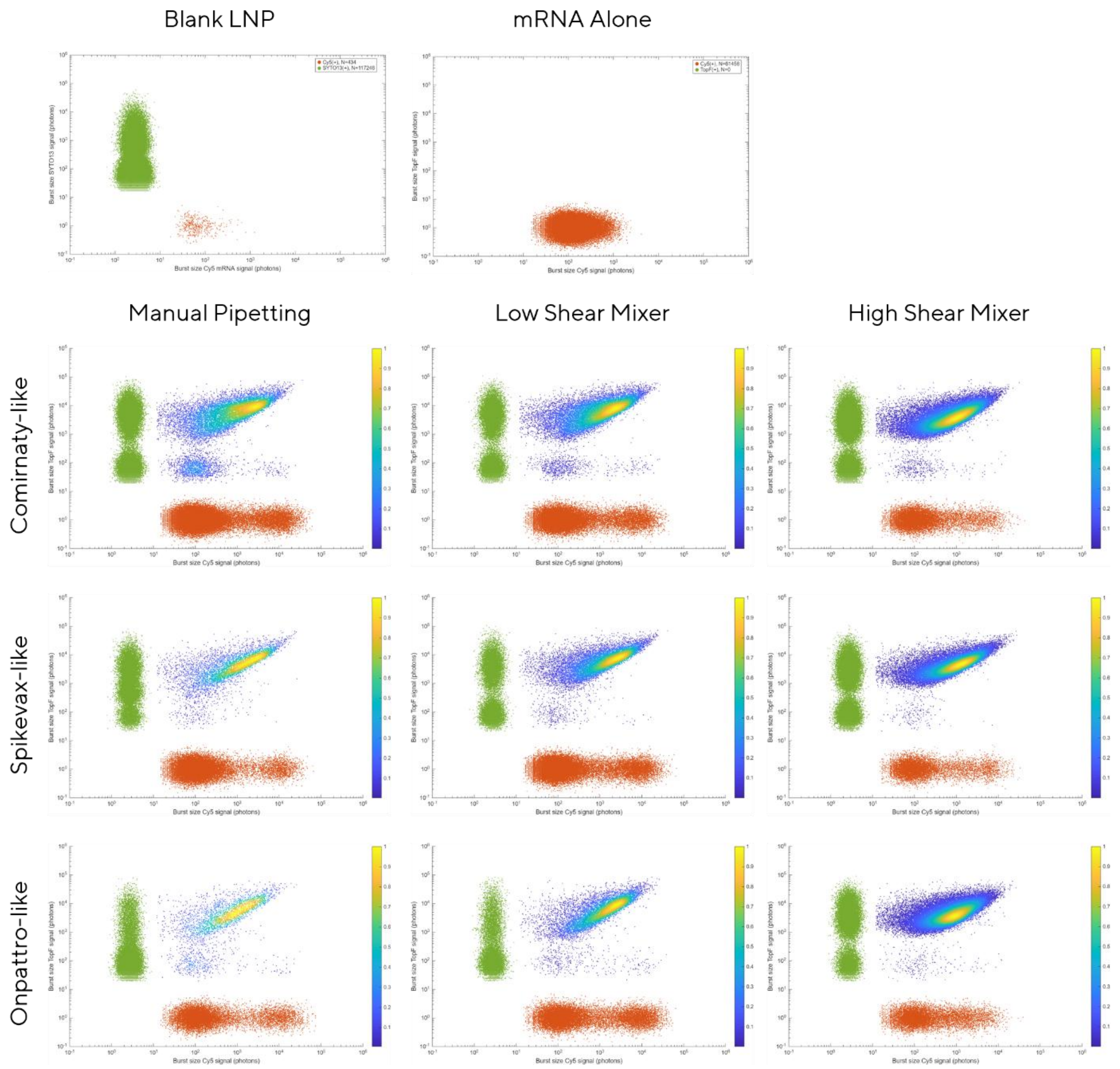


Fig. S5. Scatter plots obtained by CICS analysis.

Table S2. SPARTA analysis of particle-level Raman shift distributions of Comirnaty-like LNPs. ALC-0315 (1655–1677 cm$^{-1}$)

| | High-Shear Mixer | Low-Shear Mixer | Manual Pipetting |
|---|---|---|---|
| **N (particles)** | 245 | 248 | 246 |
| **Median** | 0.0399 | 0.0381 | 0.0291 |
| **IQR** | 0.0070 | 0.0132 | 0.0129 |
| **CQV** | 0.0873 | 0.1699 | 0.2064 |

Table S3. SPARTA analysis of particle-level Raman shift distributions of Comirnaty-like LNPs.

Cholesterol (696–714 cm$^{-1}$)

| | High-Shear Mixer | Low-Shear Mixer | Manual Pipetting |
|---|---|---|---|
| **N (particles)** | 245 | 248 | 246 |
| **Median** | 0.0127 | 0.0108 | 0.0108 |
| **IQR** | 0.0023 | 0.0026 | 0.0035 |
| **CQV** | 0.0910 | 0.1212 | 0.1690 |

Table S4. SPARTA analysis of particle-level Raman shift distributions of Comirnaty-like LNPs.

Total lipid (1415–1500 cm$^{-1}$)

| | High-Shear Mixer | Low-Shear Mixer | Manual Pipetting |
|---|---|---|---|
| **N (particles)** | 245 | 248 | 246 |
| **Median** | 0.3654 | 0.3643 | 0.3988 |
| **IQR** | 0.0336 | 0.0513 | 0.0407 |
| **CQV** | 0.0461 | 0.0703 | 0.052 |

Table S5. SPARTA analysis of particle-level Raman shift distributions of Spikevax-like LNPs.

SM102 (1120–1145 $cm^{-1}$)

| | High-Shear Mixer | Low-Shear Mixer | Manual Pipetting |
|---|---|---|---|
| **N (particles)** | 237 | 248 | 241 |
| **Median** | 0.0148 | 0.0207 | 0.0192 |
| **IQR** | 0.0046 | 0.0112 | 0.0049 |
| **CQV** | 0.1602 | 0.2884 | 0.1317 |

Table S6. SPARTA analysis of particle-level Raman shift distributions of Spikevax-like LNPs.

Cholesterol (696–714 $cm^{-1}$)

| | High-Shear Mixer | Low-Shear Mixer | Manual Pipetting |
|---|---|---|---|
| **N (particles)** | 237 | 248 | 241 |
| **Median** | 0.0126 | 0.0096 | 0.0110 |
| **IQR** | 0.0047 | 0.0038 | 0.0030 |
| **CQV** | 0.1814 | 0.1993 | 0.1348 |

Table S7. SPARTA analysis of particle-level Raman shift distributions of Spikevax-like LNPs.

Total lipid (1415–1500 $cm^{-1}$)

| | High-Shear Mixer | Low-Shear Mixer | Manual Pipetting |
|---|---|---|---|
| **N (particles)** | 237 | 248 | 241 |
| **Median** | 0.2643 | 0.3358 | 0.3613 |
| **IQR** | 0.0866 | 0.0573 | 0.0533 |
| **CQV** | 0.1573 | 0.0858 | 0.0750 |

Table S8. SPARTA analysis of particle-level Raman shift distributions of Onpattro-like LNPs. MC3 (1639–1673 $cm^{-1}$)

| | High-Shear Mixer | Low-Shear Mixer | Manual Pipetting |
|---|---|---|---|
| **N (particles)** | 250 | 248 | 246 |
| **Median** | 0.1242 | 0.1259 | 0.1220 |
| **IQR** | 0.0085 | 0.0069 | 0.0126 |
| **CQV** | 0.0341 | 0.0272 | 0.0519 |

Table S9. SPARTA analysis of particle-level Raman shift distributions of Onpattro-like LNPs. Cholesterol (696–714 $cm^{-1}$)

| | High-Shear Mixer | Low-Shear Mixer | Manual Pipetting |
|---|---|---|---|
| **N (particles)** | 250 | 248 | 246 |
| **Median** | 0.0124 | 0.0102 | 0.0118 |
| **IQR** | 0.0017 | 0.0019 | 0.0025 |
| **CQV** | 0.0668 | 0.0944 | 0.1063 |

Table S10. SPARTA analysis of particle-level Raman shift distributions of Onpattro-like LNPs. Total lipid (1415–1500 $cm^{-1}$)

| | High-Shear Mixer | Low-Shear Mixer | Manual Pipetting |
|---|---|---|---|
| **N (particles)** | 250 | 248 | 246 |
| **Median** | 0.3132 | 0.3281 | 0.3084 |
| **IQR** | 0.0246 | 0.0165 | 0.0477 |
| **CQV** | 0.0394 | 0.0252 | 0.0777 |

Table S11. mRNA payload distribution per particle of Comirnaty-like LNPs analyzed by CICS.

| | High-Shear Mixer | Low-Shear Mixer | Manual Pipetting |
|---|---|---|---|
| **Mean (copy per particle)** | 6.5 | 8.7 | 7.0 |
| **Median** | 5.1 | 7.8 | 4.6 |
| **SD** | 4.2 | 6.3 | 5.9 |
| **IQR** | 6.3 | 11.3 | 10.0 |

Table S12. mRNA payload distribution per particle of Spikevax-like LNPs analyzed by CICS.

| | High-Shear Mixer | Low-Shear Mixer | Manual Pipetting |
|---|---|---|---|
| **Mean (copy per particle)** | 7.3 | 10.4 | 8.8 |
| **Median** | 6.2 | 10.5 | 6.9 |
| **SD** | 4.6 | 6.9 | 7.4 |
| **IQR** | 7.2 | 12.7 | 13.2 |

Table S13. mRNA payload distribution per particle of Onpattro-like LNPs analyzed by CICS.

| | High-Shear Mixer | Low-Shear Mixer | Manual Pipetting |
|---|---|---|---|
| **Mean (copy per particle)** | 6.1 | 9.9 | 6.6 |
| **Median** | 5.2 | 9.5 | 3.9 |
| **SD** | 3.3 | 6.4 | 5.8 |
| **IQR** | 5.0 | 11.2 | 10.1 |

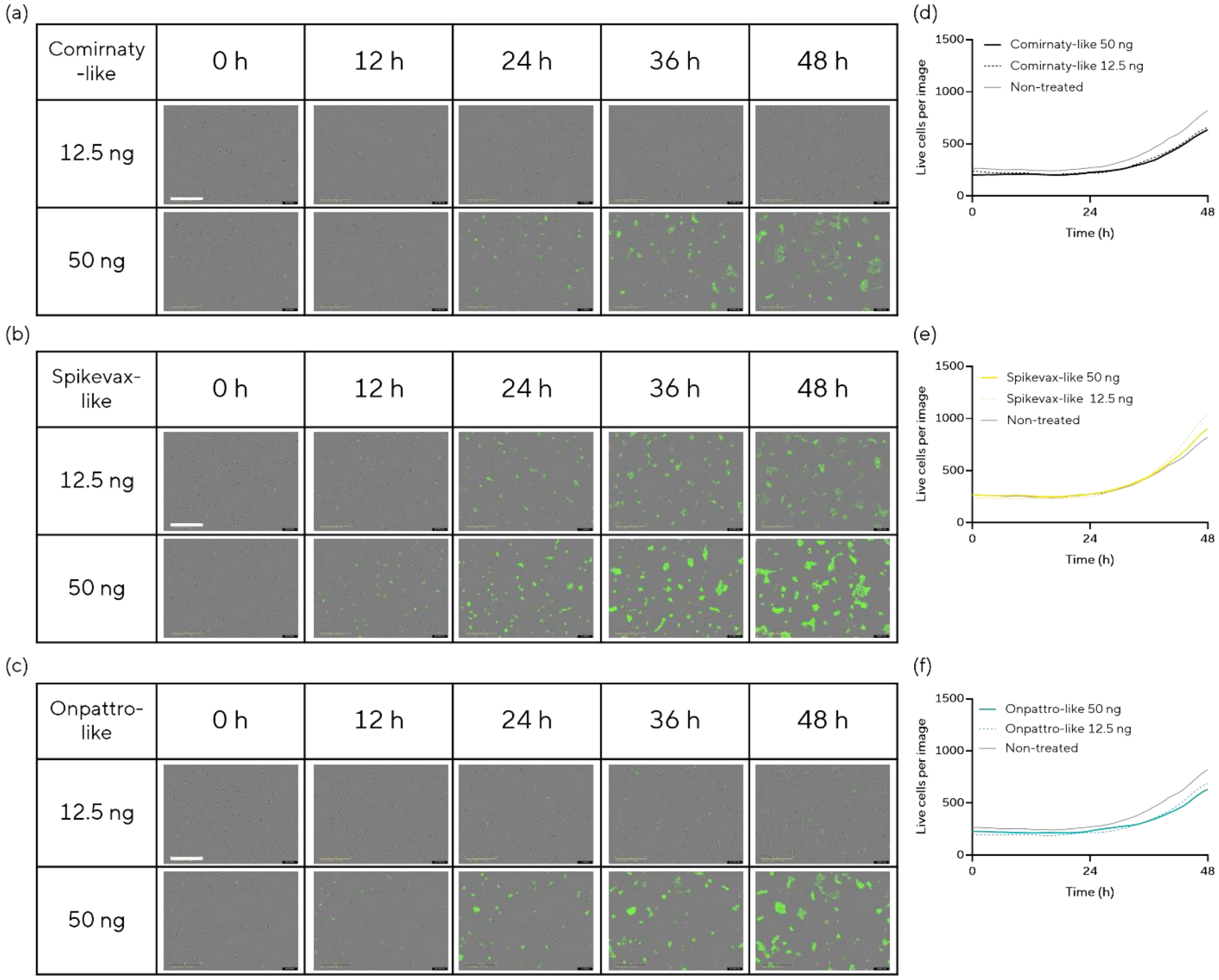


Fig. S6. *In vitro* transfection in HEK293T cells. LNPs were prepared using the high-shear mixer. Dose-dependent responses were monitored by time-lapse imaging with the IncuCyte S3. (a–c) Fluorescence images captured 0, 12, 24, 36 and 48 h after dosing. Scale bar, 400 µm: (a) Comirnaty-like, (b) Spikevax-like, and (c) Onpattro-like. (d–f) Live-cell count per image over 48 h, quantified using the IncuCyte AI Health Scan module: (d) Comirnaty-like, (e) Spikevax-like, and (f) Onpattro-like.

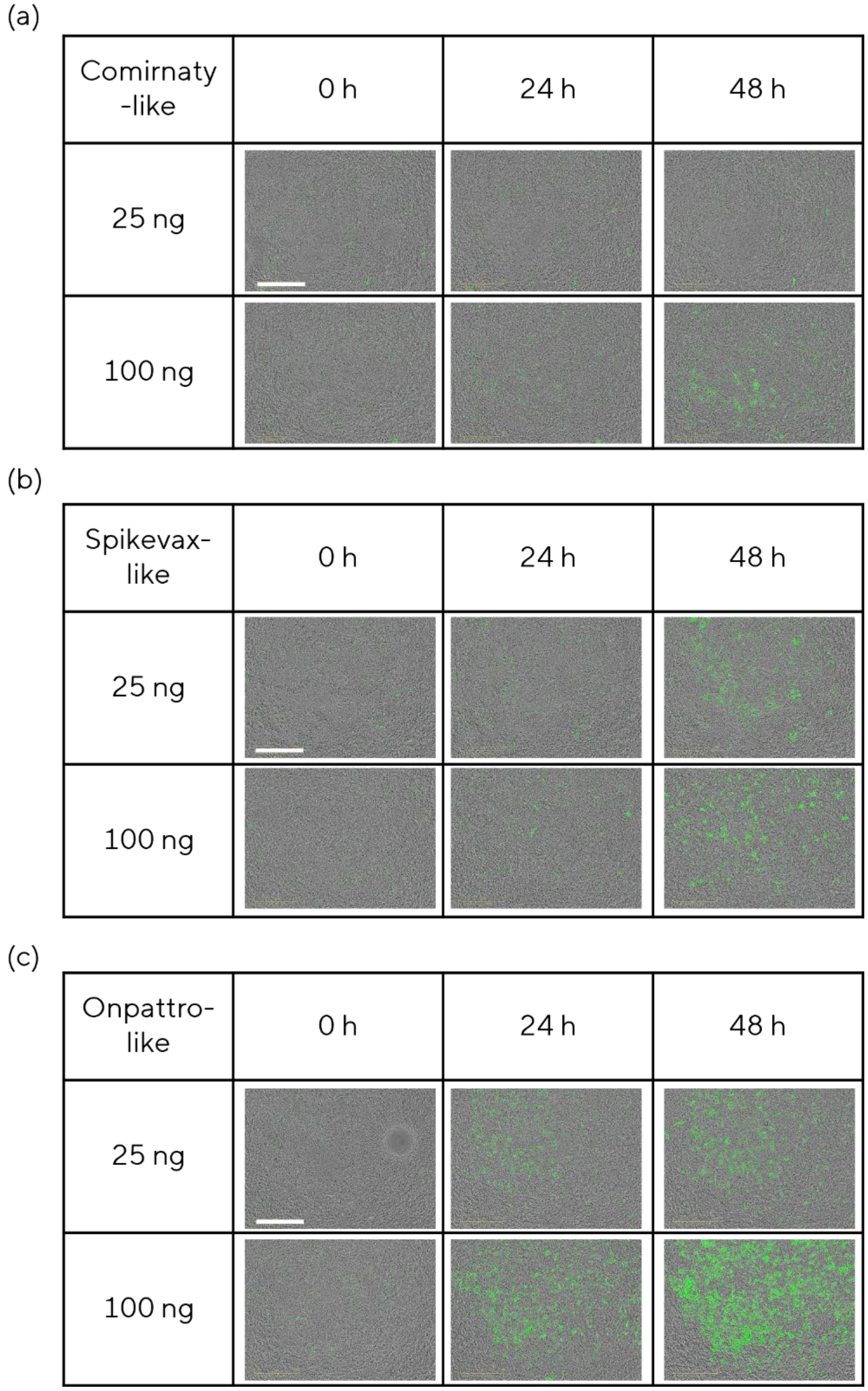


Fig. S7. *In vitro* transfection of 3D liver tissue models with lipid nanoparticles (LNPs) prepared by the high-shear mixer. Dose-dependent responses were monitored by time-lapse imaging with the IncuCyte SX5. (a) Comirnaty-like, (b) Spikevax-like, and (c) Onpattro-like. Scale bar, 400 μm.

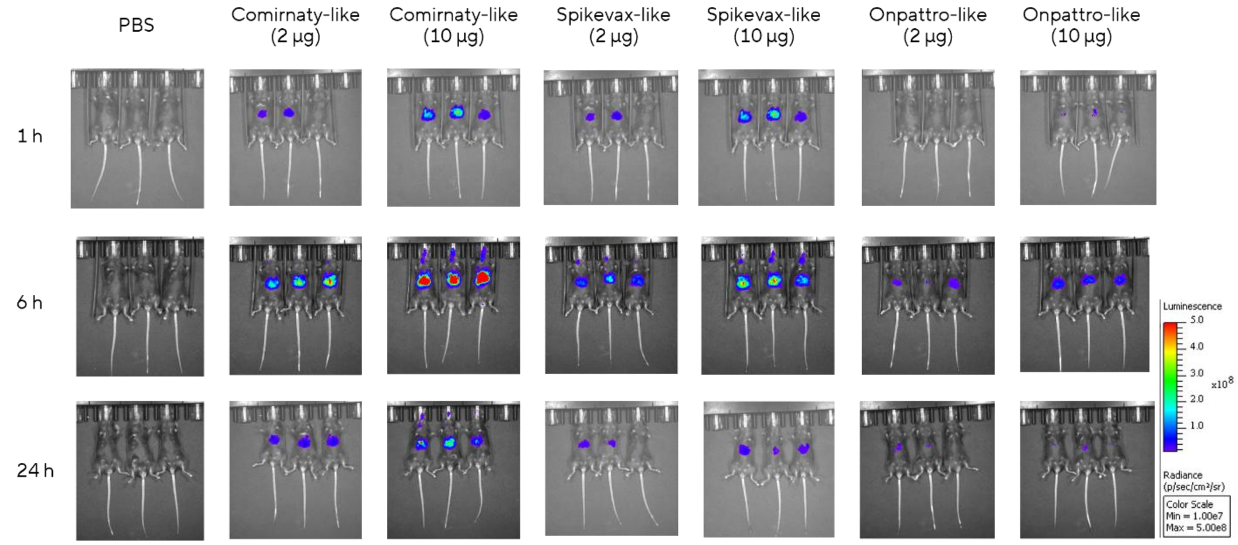


Fig. S8. Representative IVIS images of female mice treated with LNPs at each time point.

**Dimensionless Analysis of the mixing time**

Flow and transport in a spiral microchannel are characterized by the Reynolds number (Re), Peclet number (Pe), and Dean number (De): $Re = \frac{UD_H}{\nu}$, $Pe = \frac{UD_H}{D}$, $De = Re\sqrt{\frac{D_H}{R_c}}$ where U is the average flow velocity, $D_H$ is the hydraulic diameter, $D$ is the diffusion coefficient, ν is the kinematic viscosity, and $R_c$ is the radius of curvature of the channel centerline.

In curved microchannels, Dean vortices induce transverse motion that stretches and folds concentration interfaces. This process can be approximated by a stretching rate: $\lambda \sim \left(\frac{U}{D_H}\right) f(De)$ where $\frac{U}{D_H}$ represents the base shear rate and $f(De)$ describes the enhancement in stretching caused by Dean vortices.[3] The function $f(De)$ is expected to increase with Dean number. As stretching proceeds, the thickness of concentration filaments decreases exponentially:[4]

$$\delta(t) = \delta_0 exp(-\lambda t)$$

Diffusion becomes effective when the filament thickness approaches the diffusive length scale:[4,5]

$$\delta_d \sim \sqrt{\frac{D}{\lambda}}$$

Therefore, the mixing time is determined by the balance between stretching and diffusion. At the mixing time, the stretched filament thickness equals the diffusive length scale:

$$\delta(t_{mix}) = \delta_d$$

$$\delta_0 exp(-\lambda t_{mix}) = \sqrt{\frac{D}{\lambda}}$$

Taking the logarithm of both sides gives:

$$ln(\delta_0) - \lambda t_{mix} = \left(\frac{1}{2}\right) ln\left(\frac{D}{\lambda}\right)$$

Solving for the mixing time gives:

$$t_{mix} = \left(\frac{1}{\lambda}\right) [ln(\delta_0) - \left(\frac{1}{2}\right) ln\left(\frac{D}{\lambda}\right)]$$

Assuming that the initial concentration filament thickness scales with the hydraulic diameter, $\delta_0 \sim D_H$ and that the stretching rate is given by $\lambda \sim \left(\frac{U}{D_H}\right) f(De)$,

$$\delta_0/\delta_d \sim D_H \sqrt{(\frac{\lambda}{D})} \sim \sqrt{Pe\, f(De)}$$

Therefore, the mixing time can be approximated as

$$t_{mix} \sim (\frac{1}{\lambda}) ln(\sqrt{Pe\, f(De)})$$

Since constant prefactors do not affect the scaling relationship, this can be simplified to constant perfectors:

$$t_{mix} \sim \left(\frac{1}{\lambda}\right) ln\big(Pe\, f(De)\big) \sim (\frac{D_H}{U f(De)}) ln(Pe\, f(De))$$

Jiang *et al*. reported an exponential mixing coefficient, $\mu$, which is proportional to the Lyapunov stretching rate for a fixed channel geometry.[6] Therefore, we take $f(De) \propto \mu(De)$.

Over the range De∼100–200, μ is approximately proportional to De, suggesting that $f(De) \sim De$.

Substituting this approximation into the mixing time expression gives

$$t_{\mathrm{mix}} \sim \frac{D_H}{U\, De} ln(Pe\, De)$$

Thus, in this Dean-number range, the mixing time is expected to decrease approximately with increasing Dean number, while retaining only a weak logarithmic dependence on Pe and De.

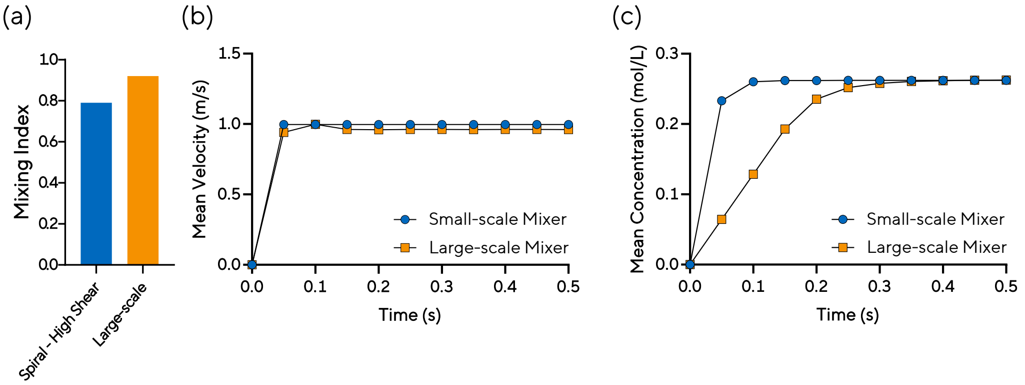


Fig. S9. CFD simulation comparing the mixing performance between the scale-up mixer and small-scale mixer: (a) mixing index at the channel outlet, (b) temporal changes in mean velocity, and (c) temporal changes in mean concentration.
